# A Structural Classification of the Variant Surface Glycoproteins of the African Trypanosome

**DOI:** 10.1101/2023.02.04.525354

**Authors:** Sara Đaković, Johan Zeelen, Anastasia Gkeka, Monica Chandra, Konstantina Foti, Monique van Straaten, Evi P. Vlachou, Francisco Aresta-Branco, Joseph P. Verdi, F. Nina Papavasiliou, C. Erec Stebbins

## Abstract

Long-term immune evasion by the African trypanosome is achieved through repetitive cycles of surface protein replacement with antigenically distinct versions of the dense Variant Surface Glycoprotein (VSG) coat. Thousands of VSG genes and pseudo-genes exist in the parasite genome that, together with genetic recombination mechanisms, allow for essentially unlimited immune escape from the host’s adaptive immune system. The diversity space of the “VSGnome” at the protein level was thought to be limited to a few related folds whose structures were determined more than 30 years ago. However, recent progress has shown that the VSGs possess significantly more architectural variation than had been appreciated. Here we combine experimental X-ray crystallography with deep-learning structural prediction using Alphafold to produce models of hundreds of VSG proteins. We classify the VSGnome into groups based on protein architecture and oligomerization state, contextualize recent bioinformatics clustering schemes, and extensively map VSG-diversity space. We demonstrate that in addition to the structural variability and post-translational modifications observed thus far, VSGs are also characterized by variations in oligomerization state and possess inherent flexibility and alternative conformations, lending additional variability to what is exposed to the immune system. Finally, these additional experimental structures and the hundreds of Alphafold predictions confirm that the molecular surfaces of the VSGs remain distinct from variant to variant, supporting the hypothesis that protein surface diversity is central to the process of antigenic variation used by this organism during infection.

## INTRODUCTION

African trypanosomiasis is a human and animal infectious disease caused by several species of protozoan parasites of the genus *Trypanosoma* [1,2]. These single-celled, eukaryotic parasites can live and reproduce extracellularly in the bloodstream of the host, and are transmitted by the tsetse fly vector (*Glossina sp*.) [3]. The geographical range of the tsetse fly in Africa correlates to the distribution of human trypanosomiasis, which today covers a region of 8 million km^2^ between 14 and 20 degrees latitude [4]. African trypanosomiasis has hampered Central Africa’s economic progress due to its impact on both human and livestock populations [5,6].

Within the blood, the trypanosome population is continuously exposed to the host’s immune system, yet is able to thrive and persist. This feat is made possible by a highly optimized system of antigenic variation, in which the ~10 million, monoallellically expressed molecules of the Variant Surface Glycoprotein (VSG) coat undergo repeated cycles of “switching”, a process by which antigenically distinct VSGs are expressed at different times [7–9]. This creates a cyclic process of trypanosome growth to high parasitemia, immune response and clearance of the dominant VSG variants, and the subsequent growth of immune-escape variants expressing different VSGs. This process renders the host in a perpetual state of infection that cannot be cleared without pharmacological intervention, typically leading to long-term morbidity and mortality [10]. Central to this process are the thousands of VSG genes and pseudogenes in the parasite genome that serve as the parasite’s extensive antigen repertoire..

The mature VSG proteins are attached to the cell surface by a GPI-anchor, and consist of two main regions: (1) a large, N-terminal domain (NTD) of roughly 300-400 amino acids that is most distal to the membrane and (2) a smaller, membrane-proximal C-terminal domain (CTD) spanning 80-120 amino acids beneath the NTD that harbors the GPI-anchor [11]. Multiple studies have shown that the CTD is minimally immunogenic [12] and quite possibly inaccessible to the immune system when it is part of the coat [12,13], which correlates well with it being much more highly conserved in sequence than the NTD [14,15]. The NTD is therefore presumed to be the most antigenic portion of the VSG. Consistent with this hypothesis, the NTD typically shows approximately 10-30% identity from variant to variant [14]. Most of the conserved residues occur in the architecturally common regions of the VSGs, whereas the molecular surfaces (as visualized by protein structures) show very little similarity [11].

Therefore, at the heart of antigenic variation in the African trypanosome are the NTDs of the VSGs. These large subdomains are elongated folds centered on a three-helix bundle scaffold. The three-helix bundle is a ubiquitous fold harbored in proteins with diverse activities, where the structural elements specific for differing functions are often coded by sequences inserted between the helices or at the termini of the scaffold. All VSGs studied to date are so structured, possessing subdomains at spatially opposite ends of the bundle: a top lobe and bottom lobe. The recent elucidation of additional VSG structures upturned the notion that all VSGs would possess highly similar protein folds to the initial structures determined in the 1990s. What is now evident from recent work is that the VSGs possess much more architectural and topological variation than had initially been appreciated [16–18].

Furthermore, for the thirty years after the determination of the first VSG structures, it was assumed that all VSGs were homodimers. However, more recent structures and biochemical analyses have shown that two broad VSG classes can be distinguished by their oligomeric state: one class harboring exclusively dimers and the other class characterized by concentration-dependent trimers (existing in solution in either a stable monomeric or trimeric state, depending on the protein concentration [16,17,19]). At the extreme densities of packing in the two-dimensional trypanosome surface coat, it is possible that all VSGs of this class will exist in the trimeric state, although no high-resolution imaging of the parasite coat that could establish this has been reported.

The uniqueness of each VSG molecular surface led to the assumption that antigenic variation occurred exclusively through amino acid sequence divergence. Recent structures and mass spectrometric analyses of VSGs have shown that this is not solely the case. Many VSGs can be modified by post-translational *O*-linked glycosylation at the top of the molecule and this modification has been shown to be potently immunomodulatory [17]. Intriguingly, the same class that adopts the trimeric oligomerization is to-date the only class in which such O-linked glycosylation has been observed.

Thus, in the last few years, many notions regarding the VSGs have had to be reexamined. Concurrently, the diversity space of the VSGs has been broadened dramatically, raising the question of whether the thousands of possible VSG proteins in the genome could be organized into a coherent schema relating sequence, structure, and function. Along those lines, several groups have undertaken bioinformatic analysis of the “VSGnome” (the set of possible VSG proteins from the trypanosome genome). Most prominent have been two papers that have used sequence clustering algorithms to divide the VSGs into subclasses [20,21]. While these papers differed in aspects of methodology and resultant classification, there was broad agreement on the classification of VSGs that appeared to be much more highly related to each other. However, none of these efforts directly took protein architecture into account, likely due to the paucity of experimentally determined structures available at the time of the analyses.

To address this issue, we use here a collection of experimentally determined structures of VSGs (including many solved over the last few years by our group, with several unpublished structures included in this manuscript) as well as hundreds of predicted structures generated by the deep-learning system, AlphaFold, that has proven powerful at the creation of accurate three-dimension protein models from amino acid sequence alone [22,23]. The combined experimental and predicted data establish a structure-based classification scheme for the VSG proteins that is generally consistent with previous attempts to classify the VSGs, but that also places classification efforts on an architectural and functional foundation.

## MATERIALS AND METHODS

### Structural determination of VSG11

Two constructs of VSG11 NTD were used in the structural studies in this manuscript (all VSGs, unless otherwise noted, are from the strain Lister 427). The first was the wild type sequence (VSG11_WT_) that (1) crystallized as a monomer in the asymmetric unit in sodium-potassium tartrate with two structures determined: VSG11_WT_-Iodine, a crystal soaked in Na/K iodine diffracting to 1.27Å, and VSG11_WT_-Oil, a crystal diffracting to 1.23 Å resolution cryo-cooled in oil, and (2) crystallized as two monomers in the asymmetric unit (not a dimer – see Results below) in ammonium sulfate (diffracting to 1.75 Å, denoted VSG11_WT_-AS). The second construct of VSG11 was a chimeric form consisting of the VSG11_WT_ NTD connected to the VSG2 CTD, denoted VSG11_N2C_. This construct crystallized with 18 monomers of VSG11_N2C_ in an asymmetric unit comprised of six trimeric assemblies (denoted VSG11_N2C_-18mer). VSG11_WT_-18mer crystals were also obtained that diffracted to nearly 3Å resolution, but the best diffracting crystals were produced by the VSG11_N2C_ construct (2.6 Å resolution). Therefore, the presence of the VSG2 CTD was not required for the formation of the 18mer form, and since in both cases the CTDs were not present in the crystallized form, it is unlikely that the 18mer form is tied in any manner to the chimeric form.

Plasmids used to generate VSG11_WT_ and VSG11_N2C_ were described in [17]. Plasmids were first linearized by EcoRV (New England Biolabs), and then transfected into VSG2-expressing cells (2T1): 10ug of each plasmid were mixed with 100ul of cells (at a concentration of between 2.5×10^7^ and 3×10^7^) in Tb-BSF buffer (90mM Na_2_HPO_4_, pH 7.3, 5mM KCl, 0.15mM CaCl_2_, 50mM HEPES, pH7.3), using an AMAXA nucleofector (Lonza) program X-001, as previously described [24]. Blasticidin at a concentration of 100ug/ml was added after 6h and single-cell clones were obtained by serial dilutions in 24-well plates and collected after 5 days.

Clones initially screened with flow cytometry for VSG2 loss of expression using a monoclonal VSG2_WT_ antibody [25] and VSG11 gain of expression with anti-VSG11 antisera. To determine the binding of antisera to live trypanosomes, 1 x 10^6^ parasites were collected and incubated with VSG2_WT_ antisera (1:4000) or VSG11_WT_ (1:1000) together with Fc block (1:200, BD Pharmingen) in cold HMI-9 without FBS for 10 min at 4°C. Cells were washed once with cold HMI-9 and resuspended in 200μl cold HMI-9 with rat anti-mouse IgM-FITC (1:500, Biolegend). After one wash with cold HMI-9, cells were resuspended in 150μl HMI-9 and immediately analyzed with FACSCalibur (BD Bioscences) and FlowJo software (v10). For a second step, clones were sequenced by isolating RNA using the RNeasy Mini Kit (Qiagen), followed by DNAse treatment with the TURBO DNA-free kit (Invitrogen) and cDNA synthesis with ProtoScript II First Strand cDNA Synthesis (New England Biolabs). The sequences were then amplified, using Phusion High-Fidelity DNA Polymerase (New England Biolabs), a forward primer binding to the spliced leader sequence and a reverse binding to the VSG 3’untranslated region. The final products were purified by gel extraction from a 1% gel with the NucleoSpin Gel and PCR clean-up kit (Macherey-Nagel) and sent for Sanger sequencing.

All VSG11 constructs were expressed in *T. b. brucei* cultured at 37°C and 5% CO_2_ in HMI-9 media (PAN Biotech) supplemented with 10% fetal calf serum (Gibco), L-cysteine and β- mercaptoethanol. The cells from 3.6 liter culture were pelleted and the VSG11 proteins purified through modifications of previously published protocols [26]. The cells were lysed with 40 ml 0.4 mM ZnCl_2_, after centrifugation (10.000 g, 10 min) the pellet was resuspended in 30 ml 20 mM Hepes/NaOH pH=8.0, 150 mM NaCl (42°C) and centrifuged (10.000 g, 10 min). The supernatant was passed through a 25 ml Q-sepharose Fast-flow column (GE Healthcare) equilibrated with 20 mM Hepes/NaOH pH=8.0, 150 mM NaCl. The flow through was collected and concentrated. The protein was further purified on a HiLoad 16/600 Superdex 200 pg column (GE Healthcare) equilibrated in 10 mM HEPES/NaOH pH=8.0, 150 mM NaCl. The fractions containing the VSG protein were pooled and concentrated.

VSG11_WT_ crystals containing only the N-terminal domain appeared after several days at 22°C using hanging drop method with 6 mg/ml VSG11 (full-length) protein in a 1:1 volume ratio against 100 mM Tris/HCl pH=7.5 1.6-1.75M sodium-potassium tartrate. CryoOil (MiTeGen) was used as cryoprotectant and the VSG11_WT_-Oil crystals flash-cooled in liquid nitrogen. The loss of the CTD is presumed to have occurred during the crystallization stage. VSG11_WT_-Iodine crystals soaked in 200 mM KI, 100 mM Tris/HCl pH 8.0 and 1.7 M NaKTartrate were flash-cooled directly in liquid nitrogen.

VSG11_WT_-AS crystals were grown at 22°C by vapor diffusion using hanging drops with a 1:1 volume ratio of 6mg/ml protein to an equilibration buffer consisting of 0.1M sodium acetate pH 4.5 and 2M ammonium sulfate. For cryoprotection the crystals were transferred to the same buffer as that used for equilibration but supplemented with 25% v/v glycerol and were flash-cooled in liquid nitrogen.

For VSG11_N2C_-18mer the crystals were grown at 22°C by vapor diffusion using hanging drop with a 1:1 volume ratio of 6 mg/ml protein to equilibration buffer containing 19 % (w/v) PEG 2000MME, 0.2 M NaCL 0.1 M MES pH 6.0. For cryoprotection the crystals were transferred to the same buffer as used for the equilibration with 25 % (v/v) glycerol and were flash-cooled in liguid nitrogen.

Native VSG11_WT_ datasets were collected at a wavelength of 1.0 Å at the Paul Scherrer Institut Villingen. Iodine soaked crystals was collected at 1.54 Å on a Rigaku X-ray generator and detector. The structure was solved from VSG11_WT_-Iodine by single wavelength anomalous diffraction (SAD) using SHELX [27] and HKL3000 suite [28]. The initial model was built using Arp/wARP [29] with PHENIX [30], COOT [31] and PDB_REDO [32] for model optimization and refinement. The structures of VSG11_WT_-Oil, VSG11_WT_-AS, and VSG11_N2C_-18mer were solved by molecular replacement using the refined VSG11_WT_-Iodine model with the PHASER package [33] of PHENIX, and the models optimized and refined with PHENIX-REFINE [30] and with cycles of manual model building using COOT [31]. Final model statistics are shown in Supplementary Table S1.

### Structural determination of VSG615

*T.b. brucei* expressing VSG615 [34] were cultured at 37°C and 5% CO_2_ in HMI-9 media (PAN Biotech) supplemented with 10% fetal calf serum (Gibco), L-cysteine and β- mercaptoethanol. The cells from 4L culture were pelleted and the VSG615 protein purified through modifications of previously published protocols [26]. The cells were lysed with 40 ml 0.2 mM ZnCl_2_, after centrifugation (10.000 g, 10 min) the pellet was resuspended in 30 ml 20 mM HEPES/NaOH pH=7.5, 150 mM NaCl (42°C) and centrifuged (10.000 g, 10 min). The supernatant was passed through a 25 ml Q-sepharose Fast-flow column (GE Healthcare) equilibrated with 20 mM HEPES/NaOH pH=7.5, 150 mM NaCl. The flow-through was collected and concentrated to final concentration 1 mg/ml.

To generate N-terminal domain (NTD) of VSG615, the concentrated protein was subjected to limited proteolytic digestion using trypsin. The VSG at the concentration of 1 mg/ml was mixed with trypsin (5 mg/ml) (Sigma Aldrich) at 1:50 trypsin:VSG ratio and incubated for 3 hours on ice. The reaction was terminated by adding PMSF to 1 mM final concentration. The protein was further purified (separating the NTD from the CTD) by size exclusion chromatography on a HiLoad 16/600 Superdex 200 pg column (GE Healthcare) equilibrated in 20 mM HEPES/NaOH pH=7.5, 150 mM NaCl. The fractions containing the NTD of VSG615 from size exclusion chromatography were concentrated to final concentration of 10 mg/ml in 500 μl of final volume. The lysine residues on the protein were subsequently methylated by reductive alkylation [35]. 10 μl of 1M borane dimethylamine complex (DMAB) and 20 μl of 1M formamide into the protein solution and mixed gently. The mixture was incubated for 2 hours in the dark at 4 °C with rotation and the entire process repeated. Prior to overnight incubation, 5 μl of 1M DMAB was added. To stop the reaction, 1M Tris pH 7.5 was added to bring the reaction to a final volume of 1 ml. The buffer was exchanged to 20 mM HEPES pH 7.5, 150 mM NaCl by size exclusion chromatography with Superdex 200 10/300 GL column. The fractions containing the methylated protein was further concentrated to 10 mg/ml for crystallization.

The methylated VSG615 protein was crystallized at 22 °C by vapour diffusion using hanging drops formed from mixing a 1:1 volume ratio of the protein with an equilibration buffer consisting of 23 % (w/v) PEG 4000, 100 mM sodium cacodylate pH=6.0, 10 mM ZnCl_2_. For data collection, crystals were soaked in the same buffer augmented to 20 % glycerol, flash-cooled in liquid nitrogen. A native dataset was collected at the Paul Scherrer Institut, Villingen. The structure was solved with molecular replacement using a model of a VSG615 trimer predicted by AlphaFold using the PHENIX package (loop regions with a pLDDT below 50 were removed for the search model).

### Structural Determination of VSG558

VSG558 was amplified from genomic DNA of *T. brucei brucei* strain Lister 427 VSG2 expressing cells. The plasmid used to generate VSG558-expressing trypanosomes was a modification of previous plasmids designed for integration into the trypanosome genome [25]. The plasmid was first linearized by EcoRV (New England Biolabs), and then transfected into VSG2-expressing cells (2T1): 10ug of plasmid was mixed with 100ul of 4×10^7^ cells in Tb-BSF buffer (90mM Na2HPO4, pH 7.3, 5mM KCl, 0.15mM CaCl_2_, 50mM HEPES, pH7.3), using an AMAXA nucleofector (Lonza) program X-001, as previously described [24]. After 6h hygromycin was added to a concentration of 25 ug/ml and single-cell clones were obtained by serial dilutions in 24-well plates and collected after 5 days.

Clones were initially screened with flow cytometry for VSG2 loss of expression using a monoclonal VSG2_WT_ antibody [25]. To determine the binding of antisera to live trypanosomes, 2 x 10^6^ parasites were collected and incubated with 200ul FITC conjugated VSG2_WT_ antisera (FITC conjugation kit, Abcam ab102884) (1:200) in cold HMI-9 without FBS for 10 min on ice. Cells were washed twice with cold HMI-9, resuspended in 200μl cold HMI-9 and immediately analyzed with a Guava EasyCyte 4HT Flow Cytometer (Luminex). For a second step, negative clones were sequenced by isolating RNA using the RNeasy Mini Kit with on-column DNAse digestion (Qiagen) and cDNA synthesis with Superscript IV first strand synthesis system (Thermo Fisher). The sequences were then amplified, using Phusion High-Fidelity DNA Polymerase (New England Biolabs), a forward primer binding to the spliced leader sequence and a reverse binding to the VSG 3’ untranslated region. The final products were purified by gel extraction from a 1% gel with the NucleoSpin Gel and PCR clean-up kit (Macherey-Nagel) and verified by Sanger sequencing.

VSG558 was expressed and purified from trypanosomes in the same manner as VSG11. After purification the full-length protein was concentrated to 6.2 mg/ml and was crystallized at 23 °C by vapour diffusion using hanging drops formed from mixing a 1:1 volume ratio of the protein with an equilibration buffer consisting of 100 mM Citric acid/NAOH pH=5.5 and 17.5% PEG 3350. The crystals appeared after 15 hours and the same condition supplemented with 25% PEG 400 or 25% MPD was used as a cryoprotectant for crystals flash-cooled in liquid nitrogen. Datasets diffracting to 1.96 Å were collected at a wavelength of 1.0 Å at the Paul Scherrer Institut Villingen. The structure was solved using molecular replacement using a model of a dimer predicted with Alphafold [22]. After model building and refinement using PHENIX and COOT, the final model contains 2 dimers in the asymmetric unit with an R-free of 25 %.

### Protein Structural Prediction with AlphaFold (ColabFold)

VSG sequences that generated AlphaFold predictions (generated using ColabFold [36]) were obtained from Rockefeller University (https://tryps.rockefeller.edu/). Structure prediction was performed for only the N-terminal domain sequences (spanning approximately 350 amino-acids, depending on the VSG). The structure prediction process consisted of five steps: MSA construction, template search, inference with five models, model ranking based on mean pLDDT and constrained relaxation of the predicted structures. The LocalColabFold command line interface was used with arguments to specify an input FASTA file, an output directory, and various options to for structure predictions. MSA was generated by MMseq, which searches sequences with three iterations against the consensus sequences of the UniRef30 [37], a clustered version of the UniRef100 [38]. The number of final MSA hits was limited using the HHblits (v3.3.0) diversity filtering algorithm [39] implemented in MMseqs2 in multiple stages (no cluster pair has a higher maximum sequence identity than 95%; enabled --qsc 0.8 threshold and disabled all other thresholds; HHblits filtering algorithm filtered within a given sequence identity bucket such that it did not eliminate redundancy across filter buckets). Template information was gathered while AlphaFold searched via HHsearch through a clustered version of the PDB (PDB70, [39]) to find the 20 top ranked templates. Improvement of the structural models was achieved through recycling three times (the software default). At the end, AlphaFold created five models, ranked by pLDDT scores.

## RESULTS

### Additional Experimental Structures to Map VSG Diversity

Three novel VSG structures published between 2018 and 2021 — VSG3, VSG13, and VSGsur [17,18] — have demonstrated that there is far more diversity in the VSG coat proteins in fold, oligomerization, topology, and post-translational modification than was appreciated. To further map the diversity space of the VSGnome, we have undertaken X-ray crystallographic studies of numerous VSGs. Guided by recent bioinformatic analysis [20,21], we sought to examine VSGs from different clustering classes. In a separate paper [16], we determined structures of three metacyclic VSGs: VSG397, VSG531, and VSG1954, representing three clustering classes. In this paper we present experimental structures of VSG11, VSG615, and VSG558.

VSG11 and VSG615 are related to VSG3 (the topologically-similar class of monomer/trimers with *O*-linked glycan PMTs), denoted as class B [40] or N4 [21] in the literature. These were examined to confirm the fold and oligomerization of this class as well as the presence of *O*-linked glycosylation. Previously determined structures of VSG3 [17] and VSG1954 [16] showed that both possess highly similar trimers in the crystals. Further confirmation of the trimeric oligomerization state of members of this class came from a biochemical analysis of several VSGs that showed evidence of concentration dependent monomeric and trimeric forms of VSG9 in solution [19]. However, unlike VSG3, the structure of VSG1954 showed no evidence of *O*-linked glycans, despite possessing a conserved glycosylation sequence [16,17]. Whether this absence is due to unknown regulatory processes yet to be identified, to the sugar being labile and removed during purification/crystallization as occurred for VSG3 [41], or to a difference between metacyclic and bloodstream VSGs, is unclear. Therefore, pursuing additional studies of related VSGs could aid in determining how many VSGs are so modified.

NTD structures of VSG11 and VSG615 were solved to 1.23Å and 3.2Å resolution, respectively (Methods, Figure 1A and 1B, Supplementary Figure 1, Table S1). Both reveal a fold similar to VSG3 (Figure 1C) as well as the presence of *O*-linked glycans on the top surface of the protein – one observed on VSG11 and two on VSG615, although with heterogeneity on the latter (Figure 1, A and B). Like VSG3, VSG11 and VSG615 present as monomers in solution (assessed by size-exclusion chromatography, Supplementary Figure 1). This crystal form of VSG11 (VSG11_WT_-Oil, see Methods and Supplementary Table 1) crystallizes as a monomer in the asymmetric unit of the crystal, but forms a trimer along a crystallographic three-fold axis of symmetry that is nearly identical to the crystallographic trimers seen with VSG3 and VSG1954 (Figure 2A). Interestingly, VSG615 crystallized with two trimers in the asymmetric unit, and these trimers align well with those of VSG3, VSG11, and VSG1954 (Figure 2A), providing additional support that class B trimer formation is not a crystallization artifact, but could represent the relevant, biological assembly.

**Fig. 1:**
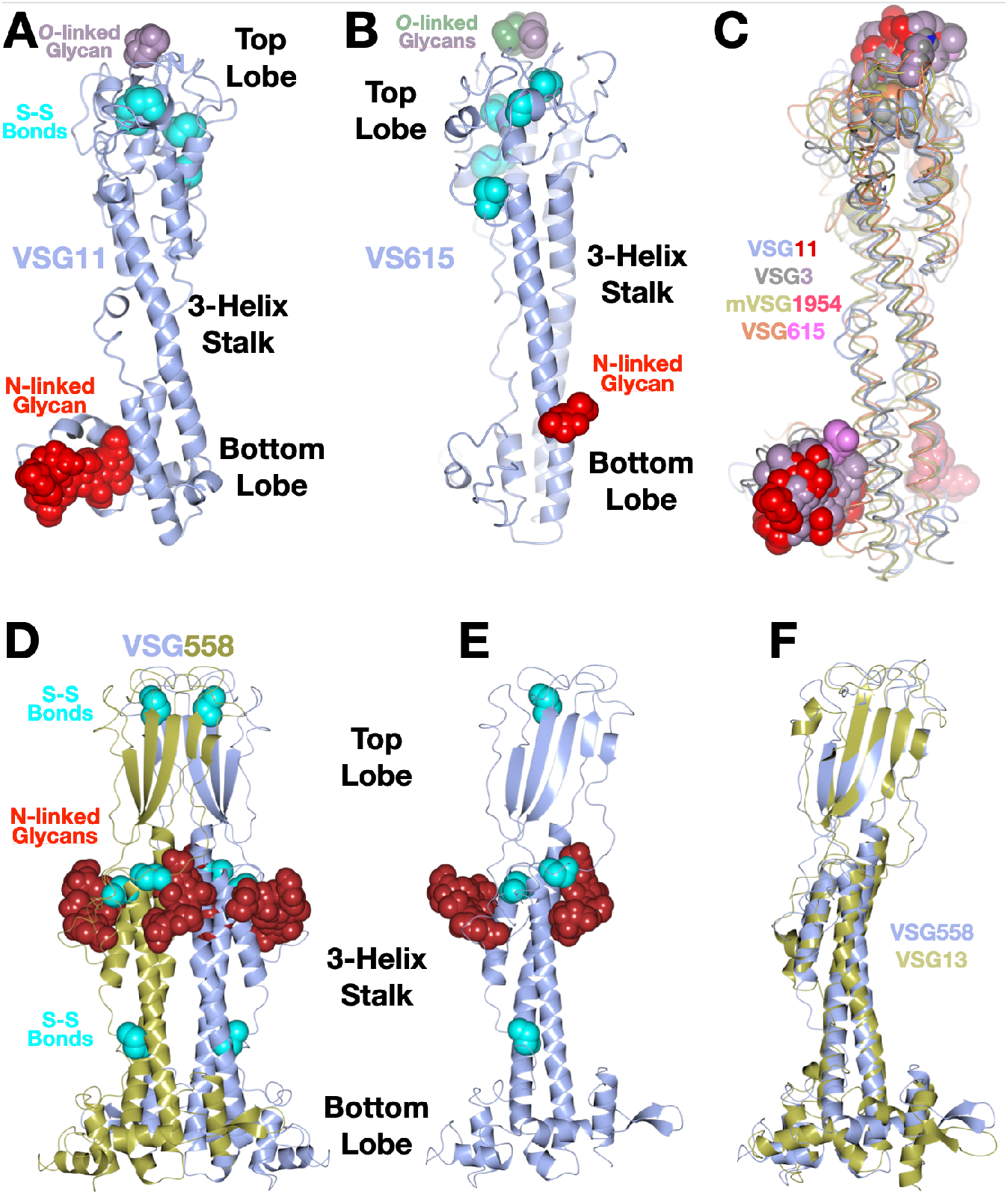
**(A)** and **(B)** Structures of the VSG11 and VSG615 NTD monomers shown as a ribbon diagram colored light blue. The N-linked glycans attached to the bottom lobe are shown in a space-filling representation colored red. Disulfide bonds (labeled S-S bonds) are shown in a space-filling representation, colored cyan. The *O*-linked glycans on the top of the molecules are shown in a space-filling representation colored purple and green (for the second sugar in VSG615). **(C)** Alignment of four B-class VSG monomers (VSG3, VSG11, VSG615, and VSG1954[16]) shown as thin “worm” drawings with each separate protein colored as indicated in the figure labels. The N- and *O*-linked sugars at the bottom and top of the molecules (respectively) are differentially colored as noted in the figure. **(D)** Structure of the VSG558 dimer illustrated as in panel (A), which each chain of the dimer colored differently. **(E)** Monomer of VSG558. **(F)** Alignment of monomers of VSG13 and VSG558. Illustrations were produced with CCP4mg [47].

**Figure 2:**
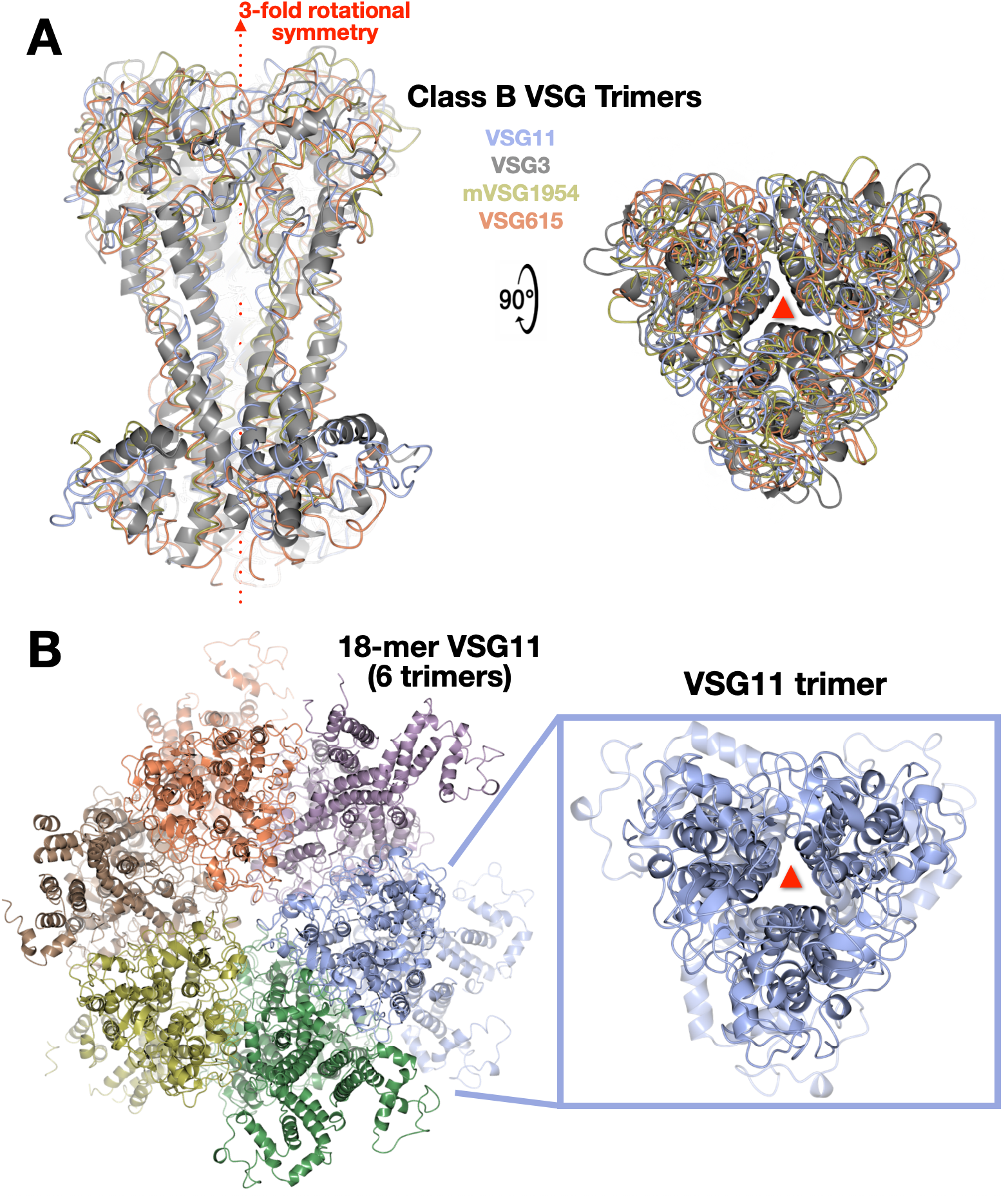
Class-B VSG Trimers. **(A)** Both crystallographic (VSG3 and VSG1954) and non-crystallographic (VSG11 and VSG615) trimeric arrangements aligned and illustrated as ribbon (gray, VSG3) or worm tracing of the mainchain (other VSGs). Two orientations are shown rotated 90 degrees. The three-fold axis of symmetry is shown in red. **(B)** The full asymmetric unit of the VSG_WT_-18mer is shown on the left with the proteins drawn as ribbon diagrams and each set of trimers colored differently. On the right is a single trimer isolated from the other six. Figures produced using CCP4mg.

An additional crystal form of VSG11 shows non-crystallographic trimerization in the asymmetric unit, while another form also raises the possibility of an inherent flexibility that could be present in the VSG proteins. In a second crystal form (VSG11_N2C_-18mer, determined to 2.6 Å resolution, Methods and Table S1), the asymmetric unit contains six trimers of the same arrangement as observed previously for all class B VSGs, for a total of 18 independent molecules (hence “18mer”, Figure 2B). In addition to helping establish that this specific trimeric arrangement is the preferred oligomerization of this class (at least at higher concentrations), the independent monomers in this asymmetric unit display a range of alternative conformations, particularly in the bottom lobe of the protein and even in some limited regions of the three-helix bundle (Figure 3A). An overall structural alignment of the monomers to themselves gives an average RMSD of 1.33 Å (RaptorX), a somewhat high value for aligning a protein to itself. Examination of the set of alignments from the monomers of the 18mer show that it appears that the divergence primarily occurs in the lower lobe (Figure 3A).

**Figure 3:**
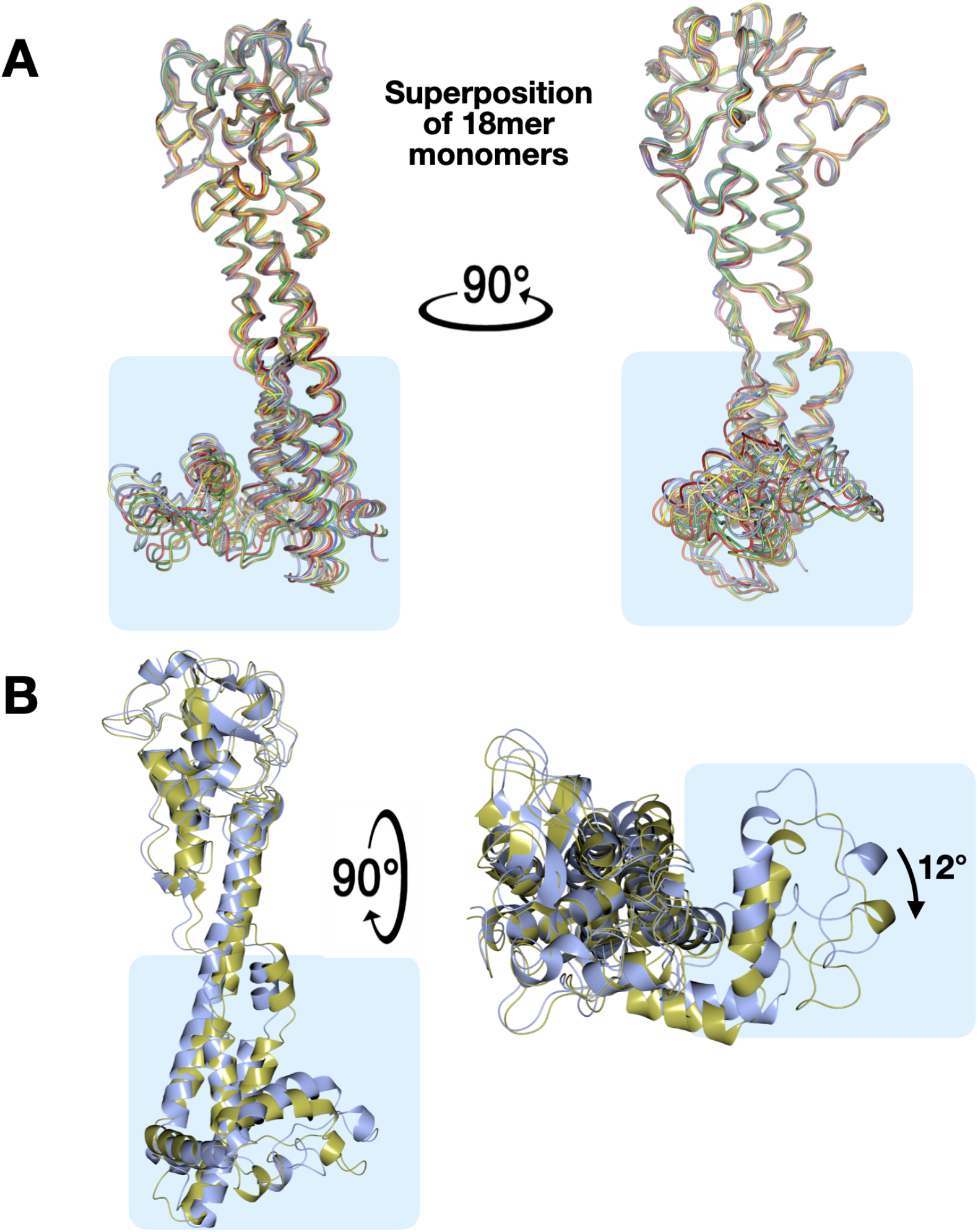
Lower Lobe Conformational Variability Observed in VSG11 Structures. **(A)**Alignment of the monomers in the asymmetric unit of VSG_WT_-18mer, each monomer given a separate color, shown in two orientations related by a 90-degree rotation about the axis of the helical bundle. **(B)** Alignment of the two monomers in the asymmetric unit of the VSG11_WT_-AS crystals. Two views are shown rotated by 90 degrees and the rotation of the bottom lobes relative to each other is indicated. Figures produced using CCP4mg.

The third crystal form (VSG11_WT_-AS grown in ammonium sulfate and solved to 1.75Å resolution, Methods, Supplementary Table S1) contains two molecules in the asymmetric unit (Supplementary Figure 2A). However, these two monomers are not arranged in a standard class A type dimeric arrangement in which the three-helix bundles, top, and bottom lobes align alongside each other, but are instead “tail-to-tail” with the lower lobe of one monomer interacting with the lower lobe of the other (Supplementary Figure 2A). In contrast, the crystal packing is characterized by a three-fold trimer identical to those seen in the all the class B protein structures solved to date, including the alternative crystal forms of VSG11 (Supplementary Figure 2B). Therefore, the assembly of VSG11_WT_-AS is most accurately described as trimeric and not dimeric (and certainly not dimers in the sense of those seen in class A VSGs), providing an interesting example in which the contents of the asymmetric unit of the crystal do not present the likely biological assembly, whereas the crystallographic arrangement does. Intriguingly, these two independent monomers are characterized by a significant conformational change in bottom lobe of the protein. This change consists of a 12-degree rotation of the bottom lobe from one monomer to the other (Figure 3B).

In addition, alignments of the VSG11_WT_-Iodine crystal form with the VSG11_WT_-Oil form (Methods and Supplementary Table S1) show that the second helix of the three-helix bundle displays a significant conformational change in the region near the middle of the bundle, the helix becoming disordered and forming a random coil in one monomer while keeping helical structure in the other (Supplementary Figure 2C). Finally, when the different monomers of the 18mer are compared with the other crystal forms described, a wide variety of conformational changes is seen, although these changes are limited to the bottom lobe and lower regions of the bundle (Supplementary Figure 3).

These alternative conformations, seen in several crystal forms where the protein is not highly constrained by the energetics of crystal packing (non-crystallographic symmetry), suggests that the VSGs on the membrane could potentially be characterized by a collection of conformers, each with molecular surfaces that differ in these regions. It is possible that such differing conformers could present distinct antigenic surfaces and impact the immunogenicity of the VSGs, combining with other forms of variability such as surface protein features and post-translational modifications to augment the diversity space of the coat.

Finally, the structure of the VSG558 NTD was determined to 1.96 Å resolution (Figure 1, D-F, Methods, Supplementary Figure 1, and Table S1). It has a highly similar structure to VSG13 [18] with a 2.54 Å RMSD over 324 residues (Figure 1F). This includes a broad and flat beta-sheet top-lobe that forms an intermolecular beta-sandwich in the dimeric assembly, a distribution of disulfides throughout the length of the rod-like structure, an N-terminus located in the “middle lobe” of the protein (instead of at the top, most membrane distal face of the VSG), and the presence of N-linked glycans directly below the top lobe. A notable difference with all the previously solved VSG NTD structures published is the presence of two N-linked glycans at the middle lobe of the VSG558 protein. To date, all VSG structures determined have been characterized by a single N-linked glycan (or none, as seen in the case of ILTat1.24). However, it has been established biochemically that some VSGs do possess more than one such sugar in the NTD, such as VSG5 [42]. The structure of VSG558 confirms the presence of multiple N-linked glycans for another VSG. This shows that not only are the locations of the N-linked glycans variable (found in both the bottom and middle lobes), but that the number of possible such carbohydrates is also variable.

### Overall classification of the VSGs based on structures

Altogether, as of the writing of this manuscript, there are twelve experimentally determined protein structures of the VSG NTD from *Trypanosoma brucei* in hand: VSG1, VSG2, VSG3, VSG11, VSG13, mVSG397, mVSG531, VSG558, VSG615, mVSG1954, VSGsur, and IlTat1.24 (“m” before the name indicating a metacyclic-stage expressed VSG). These twelve structures not only map well to the previous bioinformatic clustering schemes published previously, but they better discriminate between them and put the classification schemes on an architectural foundation. We have sought to take these structures, with an eye to the bioinformatics work, and create what we conclude is a more explanatory organizational scheme for the VSG proteins (Figure 4). To provide continuity with the previous classification schemes of Hutchinson [15] and Cross [40], we have preserved the A/B designation for classes. However, this has required a change in the meaning of subdivisions within class A to reflect insight from the protein structures (discussed below). Finally, this schema was then tested by modeling hundreds of VSG proteins with the deep learning system AlphaFold and comparing the resultant folds to the classes we created.

**Figure 4:**
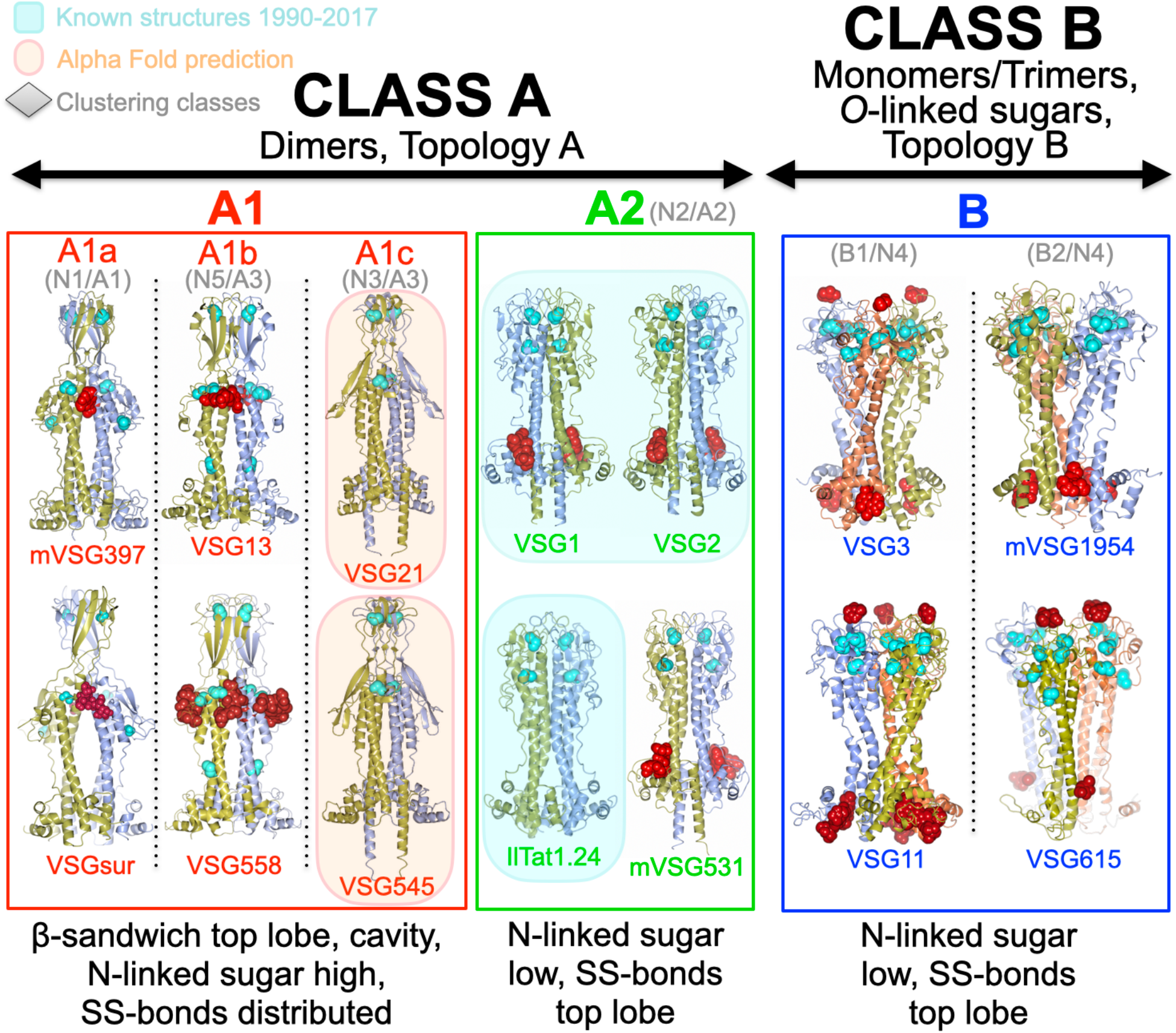
Structure-based VSG Classification Scheme. The two broad subclasses of VSG protein discussed in the main text are denoted at the top of the figure (class A and class B). Subclasses are shown in different colors with a collection of structures illustrating each class contained in a color-coordinated box matching the subclass name. Structures are drawn as in previous figures with the exception that *O*-linked glycans are colored red like N-linked glycans. Beneath each subclass name in gray are the clustering designations of the classes in previous papers discussed in the main text. Cyan backgrounds indicate structures solved prior to 2017 and orange backgrounds indicate predicted models by the AlphaFold system.

To begin, there are two broad “super-families” of VSG structures based on the topological arrangement of the bottom lobe in the primary sequence (classes A and B, Figure 4, Supplementary Figure 4). All VSG NTD structures determined have a bottom lobe subdomain structure. In class A the bottom lobe residues are present at the C-terminal portion of the NTD sequence, directly following the final helix of the bundle. In contrast, in class B the bottom lobe residues are present at the N-terminal portion of the NTD sequence between the amino acids that form the first and second helices of the bundle (Supplementary Figure 4). Further cementing this broad division into two main super-families are two observations. The first is that all class A VSGs studied to-date are found as dimers in solution and in protein crystals (the latter either as non-crystallographic dimers in the asymmetric unit or as monomers in the asymmetric unit where a crystallographic two-fold symmetry produces the dimer, [18,41,43]), whereas all studied class B VSGs are characterized by the same trimeric arrangement in the crystals (either through crystallographic or non-crystallographic symmetry, with monomers or trimers in the asymmetric unit, respectively), with biochemical evidence of monomer to trimer transitions based on protein concentration [19]. The second fact is that many of the class B VSGs are post-translationally modified by *O*-linked carbohydrates, whereas none of the class A VSGs have been found so modified. When comparing with previous efforts to classify the VSGs, our class A would correspond to class “A” in Cross [40], classes N1-N3/N5 in Weirather [21], and the older class A of Carrington [44]. Class B would correspond in these sources to classes B, N4, and B, respectively.

Within our class A superfamily are two large structural subclasses, A1 and A2 (Figure 4). In class A2 are found all the VSG structures that were solved prior to 2018 (Iltat1.24, VSG1, and VSG2), illustrating that the set of conclusions about VSG structure and function accepted for over a quarter century were based on a very limited subset of VSGs containing highly related protein folds. This subclass is characterized by a top lobe that contains all the cysteine disulfides in the NTD, a top lobe fold that is a hodge-podge of alpha helices and beta-strands, and N-linked glycan chains located in the bottom lobe of the NTD. In sharp contrast, subclass A1 VSGs are significantly longer due to the presence of a large beta-sheet subdomain for the top lobe (forming a beta-sandwich in the dimer). Further distinguishing A1 from A2 is the distributed nature of the disulfide bonds (present throughout the length of the VSG in A1), the presence of a “middle lobe” of secondary structure straddling the beginning of the three-helix bundle, and the location of the N-linked sugar just below the beta-sandwich top lobe. This dramatically different arrangement in structural elements is reflected by the differing positions of the N-terminus of the NTD, namely toward the middle of the VSG fold in A1 but located at the very top of the fold in A2. Additionally, the folds of the A1 VSGs from experimental and predicted structures (see below) suggest that this subclass should be further subdivided into three groups based on the size, conformation, and twist of the top lobe beta-sheet and the width of the space between the three-helix bundles in the dimer. These subclasses of A1 were in previous sequence clustering classification systems denoted as the separate classes A1 and A3[40] and N1, N3, and N5 [21]. However, all these subclasses are structurally similar to each other in the manners described above while differing markedly from the A2/N2 classes. Therefore, we considered it better to combine A1/A3 and N1/N3/N5 into a single class, A1 (a family contrasted to A2), and then subdivide them within A1: A1a, A1b, A1c (Figure 3).

Finally, Cross et al. [40] divide the class B VSGs into two subgroups, whereas Weirather et al. [21] do not subdivide their equivalent class, N4. In contrast to the marked structural divergences between classes A1 and A2, and even the differences within the distinct subclasses of A1, we find no broad structural differences within the class B VSGs (examining features such as the protein fold, disulfides, N- or O-linked glycans, or oligomerization). However, three more subtle differences between members of the B class can be used to divide them into two subgroups. For example, two helical regions differ between the B1 and B2 subgroups. One helix that exists in the top lobe of the B2 class is not generally present in the B1 class (Supplementary Figure 5A). Secondly, one of the bundle helices in class B1 is disordered in places relative to the same helix in many class B2 members (Supplementary Figure 5B). Thirdly, in class B2 several VSGs possess an NTD with more amino acids. In the AlphaFold predictions discussed below, these additional residues are predicted to form both an extended loop in the disordered region of the helix and also longer loops in the top lobe (Supplementary Figure 5A). Buttressing this subdivision of class B, when multiple B1 and B2 VSGs (from experimental and AlphaFold predicted models) are analyzed by structure (using the Dali Server [45]), these are divided into two classes consistent with the B1 and B2 groupings produced by sequence analysis (see the structural dendogram in Supplementary Figure 5C). We have therefore denoted a split in Figure 4 between the B1 and B2 classes.

### Mapping the VSGnome with AlphaFold

The AlphaFold deep learning artificial intelligence system is software developed by Alphabet/Google’s DeepMind program [22,23,36]. In 2020, AlphaFold established itself as a breakthrough technology with robust accuracy in the prediction of protein structure from amino acid sequence, leading many to suggest that the “protein folding problem” had been solved. We sought to utilize AlphaFold to quickly examine the sequences of hundreds of VSG proteins and compare the structural predictions both to experimentally determined VSGs as well as assess the fit of our classification scheme to the VSGnome as a whole.

We began by assessing the accuracy of AlphaFold against known VSG structures that existed in the PDB database at the time of writing this manuscript, and which would therefore be accessible to AlphaFold as structural templates. In addition, as the VSGs are multimers (dimers and trimers) and AlphaFold can predict both isolated folds and folds as oligomers [36], we generated models for monomer VSG sequences as well as for the appropriate oligomer for a given VSG sequence. Using ColabFold (Methods) and an input sequence of these VSGs in the database, we examined five ranked models output by the software from which the rank 1 model (highest pLLDT score, see Methods) was compared with experimentally solved structures. For both monomer and multimer predictions, a LocalColabFold search found the same three VSG structures to use as templates: VSG1, VSG2, and IlTat 1.24, after which the models were obtained using default settings. All VSG structures predicted by AlphaFold were of high accuracy (Figure 5A), with multimer predictions showing lower RMSD values compared to monomer predictions.

**Figure 5:**
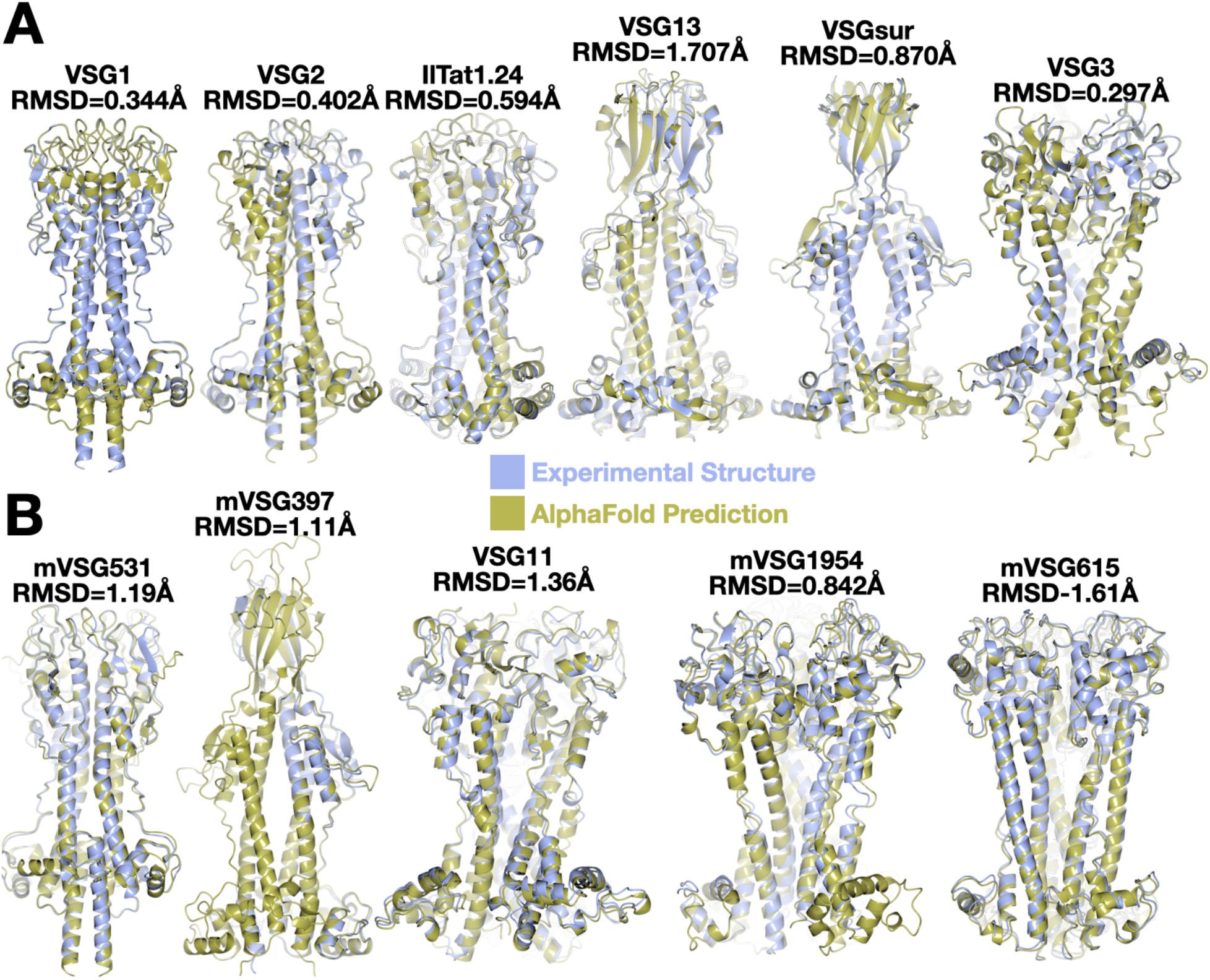
**(A)** Alignment of experimentally determined structures (blue) and AlphaFold models (gold) for structures whose coordinates were available at the time of the writing of this manuscript for use by AlphaFold as a template. **(B)** Alignment of experimentally determined structure (blue) and AlphaFold model (gold) for structures whose coordinates were not yet available at the time of the writing of this manuscript for use by AlphaFold as a template. RMSD indicates the root-mean-square deviation in the Ca positions between the two structures. Alignments and illustrations produced by CCP4mg.

Next, we repeated this analysis against structures not yet available in the protein database at the time of the analysis, namely VSG397, VSG531, and VSG1954 [16] and VSG11 and VSG615 from this manuscript. These recently determined structures were well-predicted by AlphaFold, although with slightly higher RMSD values as compared to the predictions for VSG structures already in the database (Figure 5B). Typically in these models, the three-helix bundle region showed the most accurate alignment, while discrepancies appeared mostly in loop regions connecting secondary structure. The predicted location of disulfide bonds matches very closely those observed in the experimental structures (Supplementary Figure 6). Finally, as noted in the Methods, the structures of VSG558 and VSG615 were solved by molecular replacement using the AlphaFold model of these VSGs as search models, indicating a very close fit of the model to the experimental data. These robust results from AlphaFold provide confidence that it can be used as a tool to rapidly and robustly map the structural diversity of the VSGnome.

Focusing on the Lister427 strain database (Methods), we examined 215 NTD sequences with AlphaFold and compared these predictions to the existing structures known. Our results can be summarized by stating that all of the predicted structures conform to our classification scheme and no structure was predicted that conflicted with our schema (Figure 6). We also examined 83 sequences from a second *Trypanosoma brucei* strain, *T. brucei brucei* 927. All VSGs examined from Tb 927 were predicted to adopt structures fitting our classification scheme (Supplementary Figure 7). Extending our analysis to more distally related species and VSGs like *T. vivax* showed less confidence in the modeling (Supplementary Figure 8), and therefore it is likely that template experimental structures (which currently do not exist) may be required on which the AlphaFold system can train to produce more reliable models.

**Figure 6:**
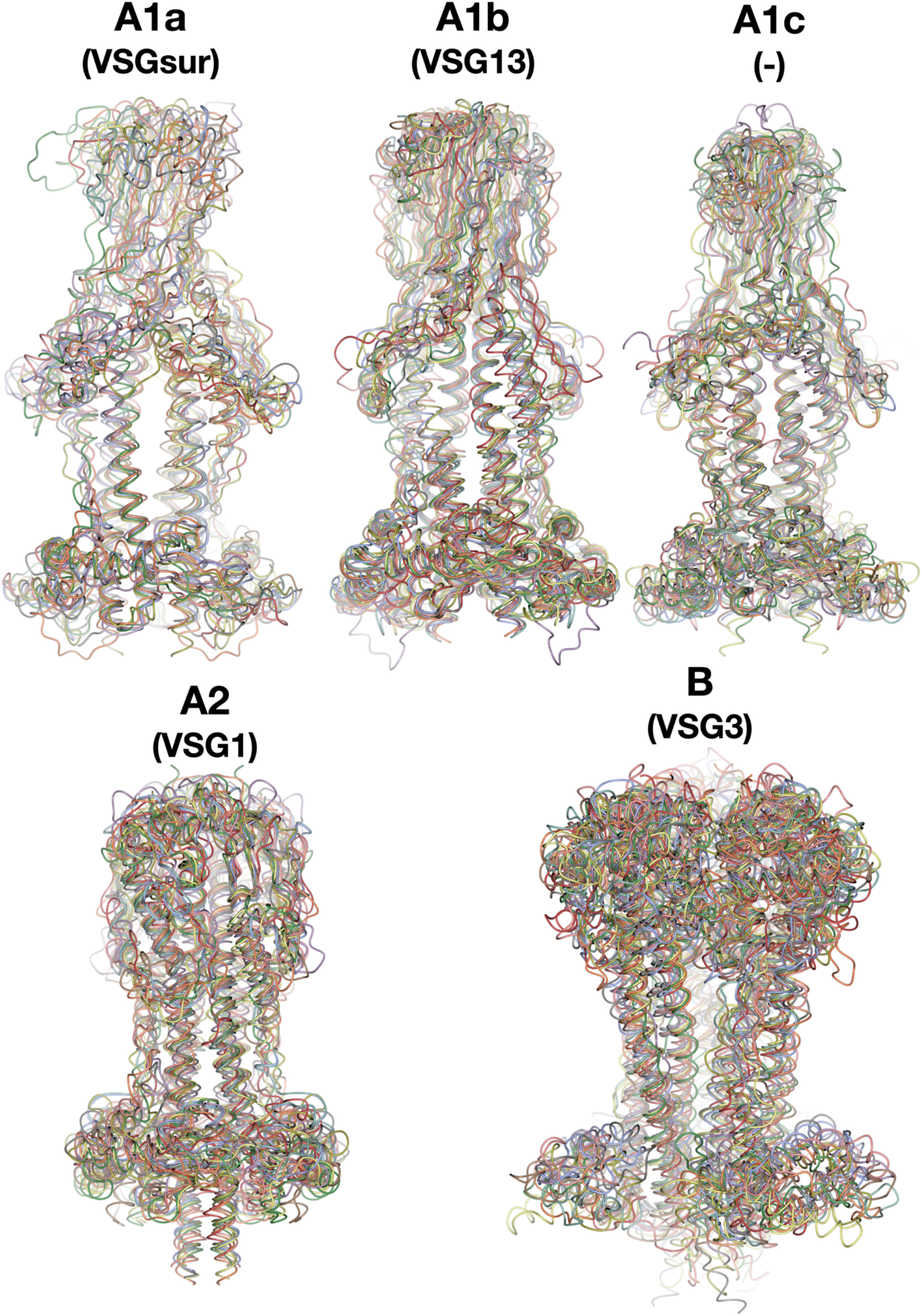
AlphaFold Models of Sequences from VSG Structural Classes. Structures are drawn in different colors with a thin worm style. **(A)** Class A1a with models for the NTD sequences of VSGsur, VSG478, VSG483, VSG485, VSG507, VSG520, VSG523, VSG529, VSG535, VSG537 **(B)** Class A1b with models for the NTD sequences of VSG13, VSG510, VSG650, VSG660, VSG661, VSG662, VSG663, VSG671, VSG717, VSG719 **(C)** Class A1c with models for the NTD sequences of VSG21, VSG391, VSG479, VSG493, VSG526, VSG545, VSG558, VSG584, VSG612, VSG659 **(D)** Class A2 with models for the NTD sequences of VSG1, VSG472, VSG476, VSG494, VSG495, VSG504, VSG514, VSG517, VSG525, VSG527, VSG536 **(E)** Class B with models for the NTD sequences of VSG3, VSG473, VSG475, VSG480, VSG486, VSG491, VSG506, VSG513, VSG516, VSG528, VSG534. Alignments generated and illustrated with CCP4mg, each structure aligned in a different color [47].

From this it can be concluded that the space of protein structure for the *T. brucei* VSGs can be well-described by the classes we have presented in this manuscript. As an overarching observation, both the experimental structures and the predicted AlphaFold models all display unique molecular surfaces in terms of the distribution of chemical properties exposed to the immune system (charge, hydrophobicity, topography, etc., Supplementary Figure 9). This is of course consistent with the notion that the VSGs are each antigenically distinct, presenting unique faces to the immune system.

## DISCUSSION

A mechanistic understanding of antigenic variation in the African trypanosome will require a complicated mixture of knowledge regarding the epitope space exposed to the host immune system and how the repertoires of antigen recognition in immune cells respond to this chemical space in the dynamic process of coat switching and the adaptive immune response [41]. At the most basic level, such an understanding necessitates a thorough knowledge of VSG antigenic diversity. Therefore, understanding the possibilities and constraints of VSG protein architecture is a critical step in this direction.

We have presented in this manuscript and in recently published work a collection of new experimental VSG structures. Combined with those already known, we have developed a structure-based classification scheme for the *T. brucei* VSGs that shows the breadth of possible protein folds for the major coat protein of the African trypanosome as well as constraining the underlying conformational possibilities inherent in the VSGnome. This classification scheme is enriched by including hundreds of VSG structural models by predicting folds using the deep learning system, AlphaFold. This revolutionary process was first validated on the VSG proteins by predicting the folds of experimentally determined VSGs in the database as well as several novel VSG structures we have determined that had not been published (and thus could not serve as templates for AlphaFold). AlphaFold predicted all with high accuracy, always performing better (as determined by match to the experimental structures) when asked to predict an oligomer of the right number for a given VSG (dimer or trimer). Since protein folding can be partially dependent on higher-order protein-protein associations, this suggests that the deep learning training of the AlphaFold algorithms has captured some aspect of the energetics of this dependency. Once we were confident in the predictions of AlphaFold for the VSG proteins, we analyzed hundreds of sequences from *T. brucei*, establishing that all predicted structures fall into the groups of our proposed classification system.

This provides an architectural framework to understand the VSGs, integrating the key structural and biochemical features of each class (protein fold and topology, oligomerization, positions of disulfides and N-linked glycans, presence of surface *O*-linked glycosylation). The importance of these constraints and the broadening of previous knowledge can be seen in the incorrect assumptions that were derived from the first three VSG structures determined over the period of 1990-2017 (VSG1, VSG2, and IlTat1.24). By happenchance, these three VSGs are not a random sampling of the VSGnome, but are clustered by structure into a single VSG class (A2) with specific oligomeric and structural properties that differ markedly from those in the other VSG classes. Our classification scheme and examination of hundreds of VSGs from multiple species suggests that the greater diversity that we have mapped out is likely close to the full space of variability in structure in *T brucei*, and likely also covers a substantial portion of the VSGs in related genomes like *T.b.* strain 927. It is quite possible that the VSGs in even distantly related species like *T. vivax* will find substantial fit with most of the *T. brucei* classifications, although currently some VSGs in *T. vivax* do not model well with AlphaFold, suggesting that there could exist novel structural classes in this and other species.

In this sense, while it was mistakenly assumed the “VSG-folding problem” was understood decades ago, it is now likely that this is indeed beginning to be realized, and that further work across the breadth of trypanosome species will be able to fully map the antigenic space of these molecules. While AlphaFold is clearly a primary choice for this structural mapping (as perhaps are newer systems like Meta AI, [46]), it has to be noted that the system currently has limitations. One caveat is that only experimental structures (or biochemical mapping) can presently establish the locations of post-translational modifications, modifications that have been shown to be immunomodulatory. Additionally, as shown with attempts to model VSG sequences from *T. vivax*,the low confidence scores of some of these predictions suggest that AlphaFold is more robust at present when there exist structural templates of sequences in specific families, e.g., *T. vivax* experimental structures are needed as templates rather than relying solely on *T. brucei* templates.

The next steps in understanding antigenic variation in the African trypanosome should likely now focus on mapping immunogenic epitopes on the VSG surfaces combined with antibody-VSG complex structures. For the time being, this will likely rest in the realm of experimental structural biology, as neither AlphaFold nor any other software has yet succeeded in robustly predicting the structures of protein-protein interactions. Once many of these have been determined, efforts to categorize the classes and natures of epitopes and the modes of interaction with antibodies will be the final piece in the puzzle to begin to develop a mechanistic framework for how this pathogen can continuously evade even the most potent immune responses.

## SUPPLEMENTARY FIGURES AND TABLES

Figures 1-9

Tables 1

## ACKNOWLEDGEMENTS

We acknowledge synchrotron time at the at the Paul Scherrer Institut, Villingen, Switzerland (SLS, beamline PX I and PX III).

## AUTHOR CONTRIBUTIONS

M.C. purified and crystallized VSG615 and J.P.Z. solved the structure. A.G. and P.V. cloned, expressed, purified, and crystallized VSG11_WT_-AS and VSG11_N/2C_-18mer. J.P.Z. and K.F. solved the structures of the VSG11_WT_ and VSG11_N/2C_-18mer. VSG11_WT_ (for the VSG11_WT_-Oil and VSG11_WT_-Iodine crystals) was purified and crystallized by J.P.Z from clones and initial conditions produced by F.A.B. M.v.S. cloned VSG558 and J.P.Z. crystallized the VSG558 and solved its structure. AlphaFold models were produced by J.P.Z. and S.D.. C.E.S. conceived of the projects, aided in structural analysis, co-wrote the manuscript, and generated figures. All authors reviewed the manuscript.

## Notes

### Competing Interest Statement

The authors have declared no competing interest.

