## Supplementary Data for "A Structural Classification of the Variant Surface Glycoproteins of the African Trypanosome"

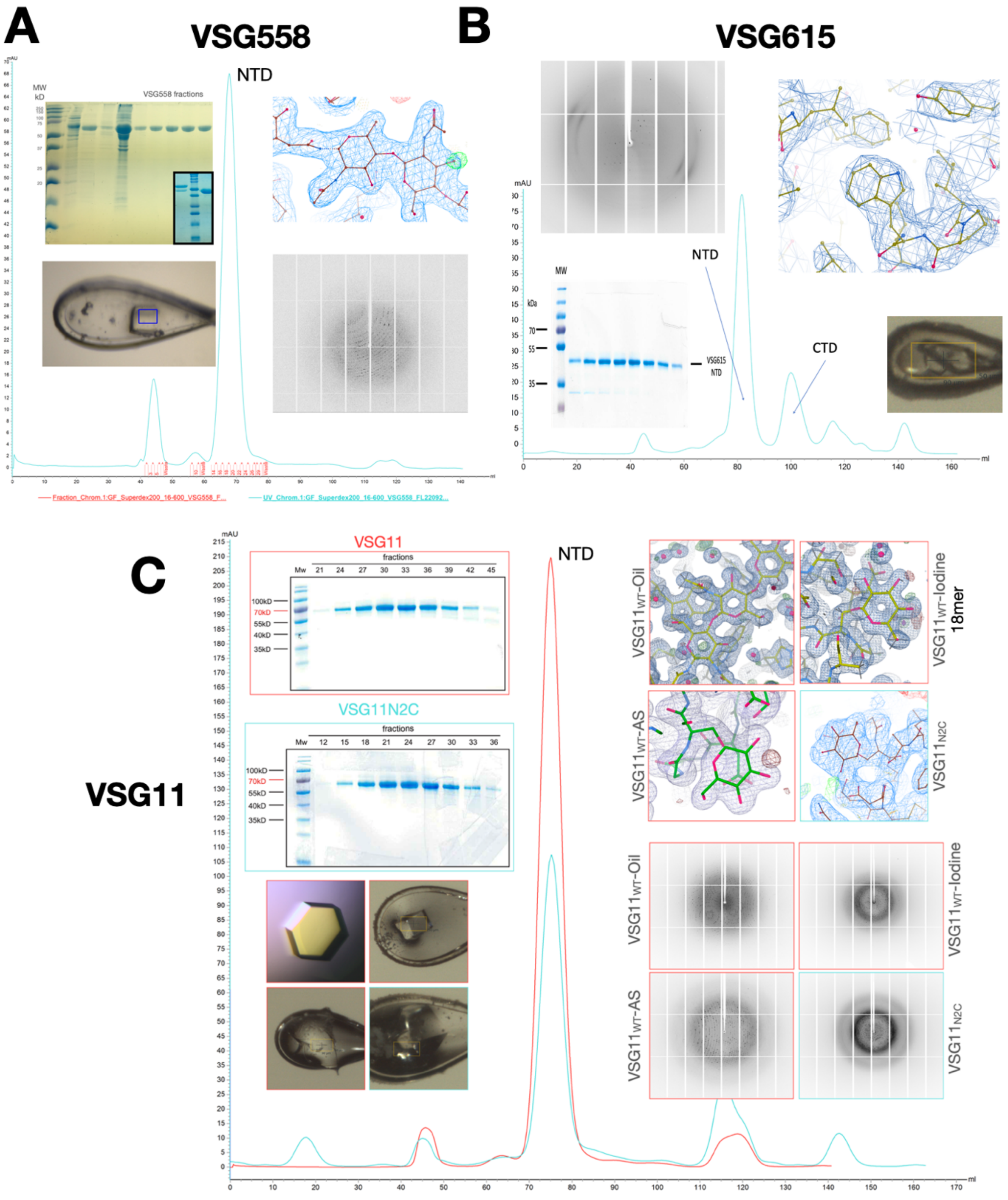


**Fig S1. Purification, Crystallization, and Representative Electron Density of VSG11 and VSG615.** Summary of various steps in the crystallographic structural solution. Panels showing the gel filtration chromatogram (Superdex 200, Methods) of purified **(A)** VSG558, **(B)** VSG615, and **(C)** VSG11 with a coomassie stained SDS-PAGE gel of the final material used for crystallization. Images of crystals grown in hanging drops, X-ray diffraction, and the final model 2Fo-Fc electron density contoured at 1σ are added alongside the chromatograph.

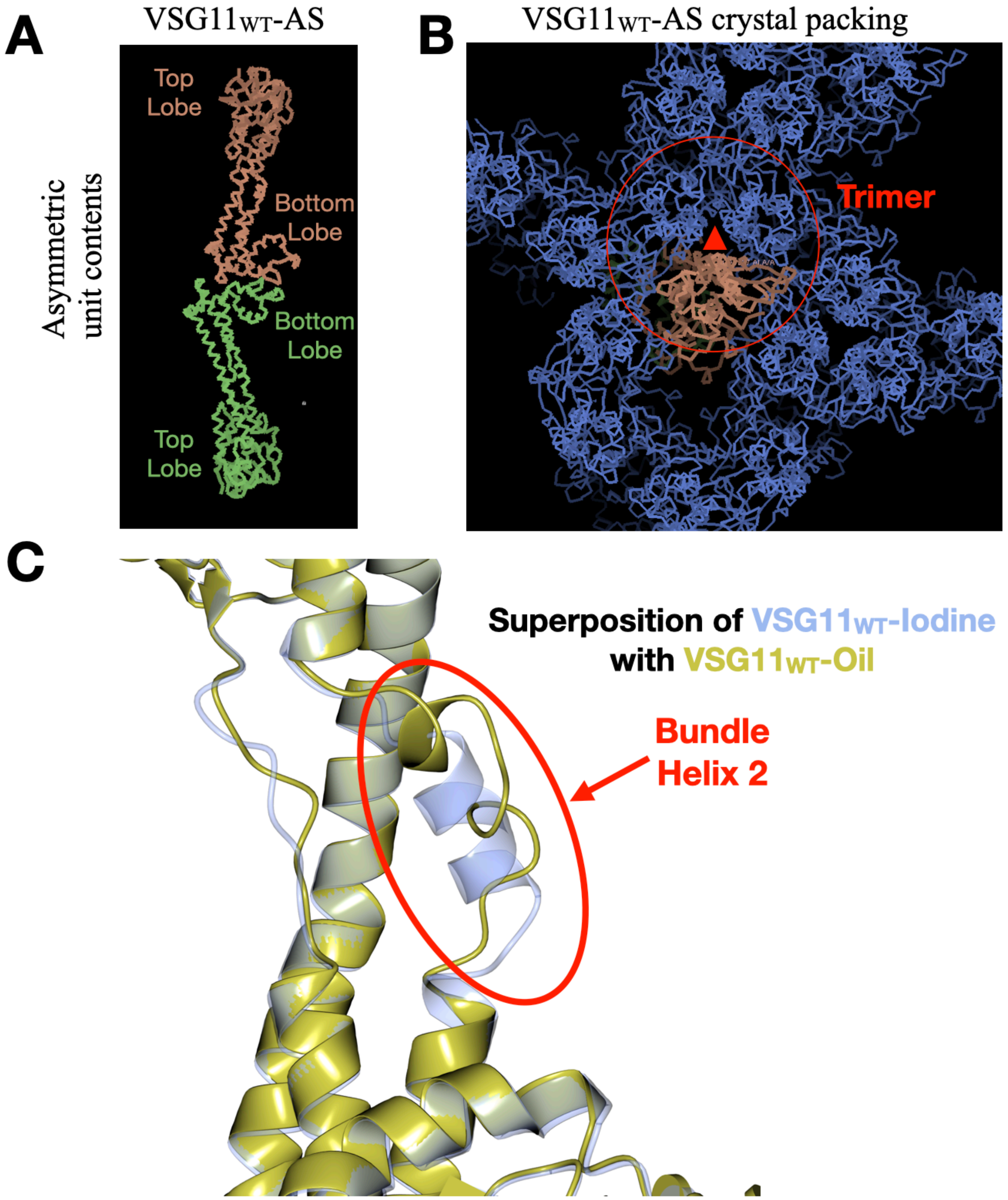


**Fig S2. Alternative Crystal Forms of VSG11. (A)** The asymmetric unit of the wild type VSG11 NTD structure from ammonium sulfate. Two molecules of VSG11 are packed "head-to-tail" and shown in green and salmon. **(B)** The full crystal packing in the form from (A) with the salmon monomer shown and the crystal-packing that produces the standard B class trimer highlighted with a red circle. **(C)** A structural alignment of two wild type VSG11 models showing the conformation change at bundle helix 2 (VSG11$_{WT}$-Iodine in blue, VSG11$_{WT}$-Oil in gold).

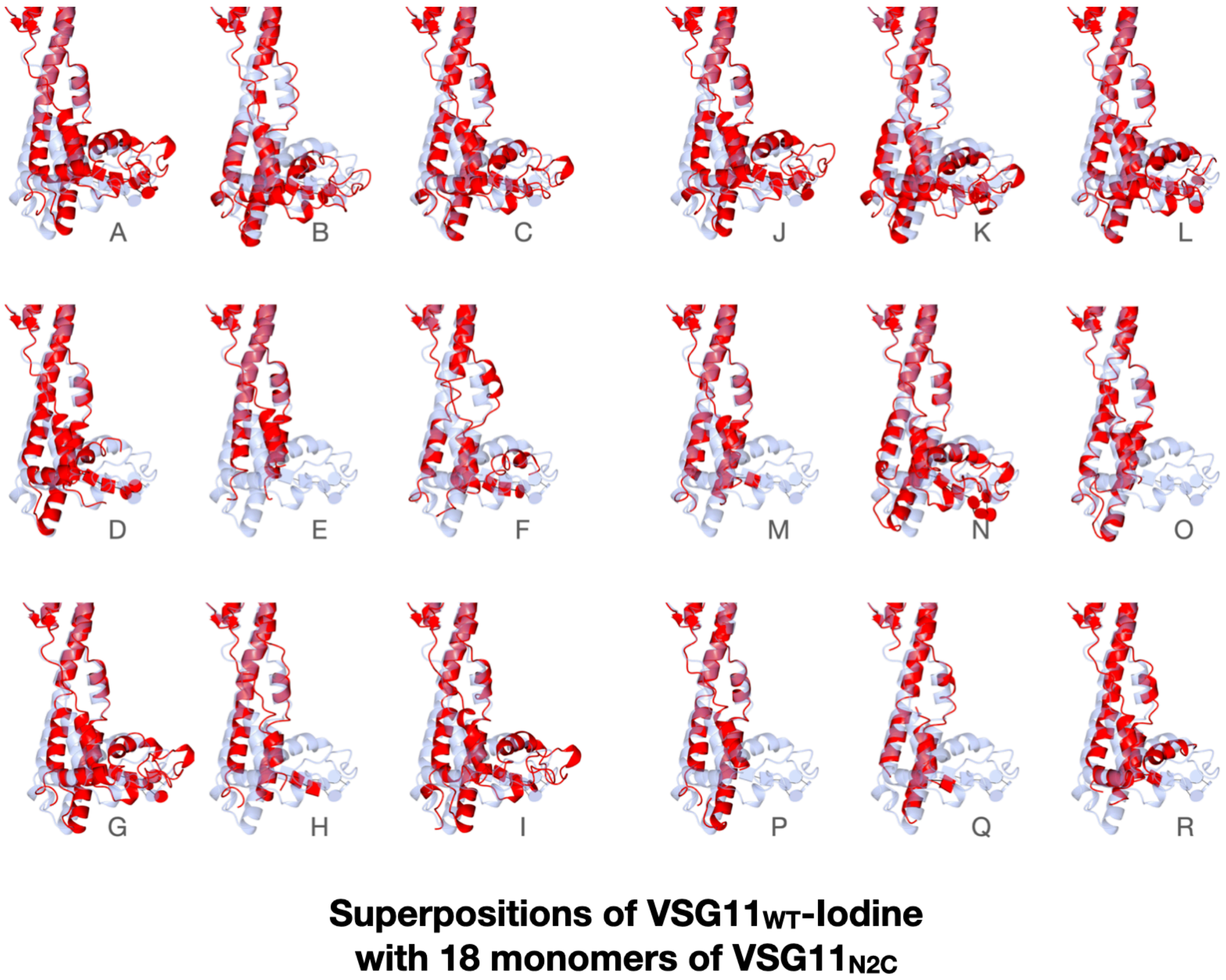


**Fig S3. Alignment of the wild type VSG11 structure soaked in iodine with the 18 monomers in the chimeric VSG11 18mer.** (A)-(R) show the $VSG11_{WT}$-Iodine in light blue and a different, individual $VSG11_{N2C}$ monomer in red. Due to the high flexibility of the bottom lobes in the 18mer crystals, not all amino acids in some monomers of the bottom lobe could be modeled.

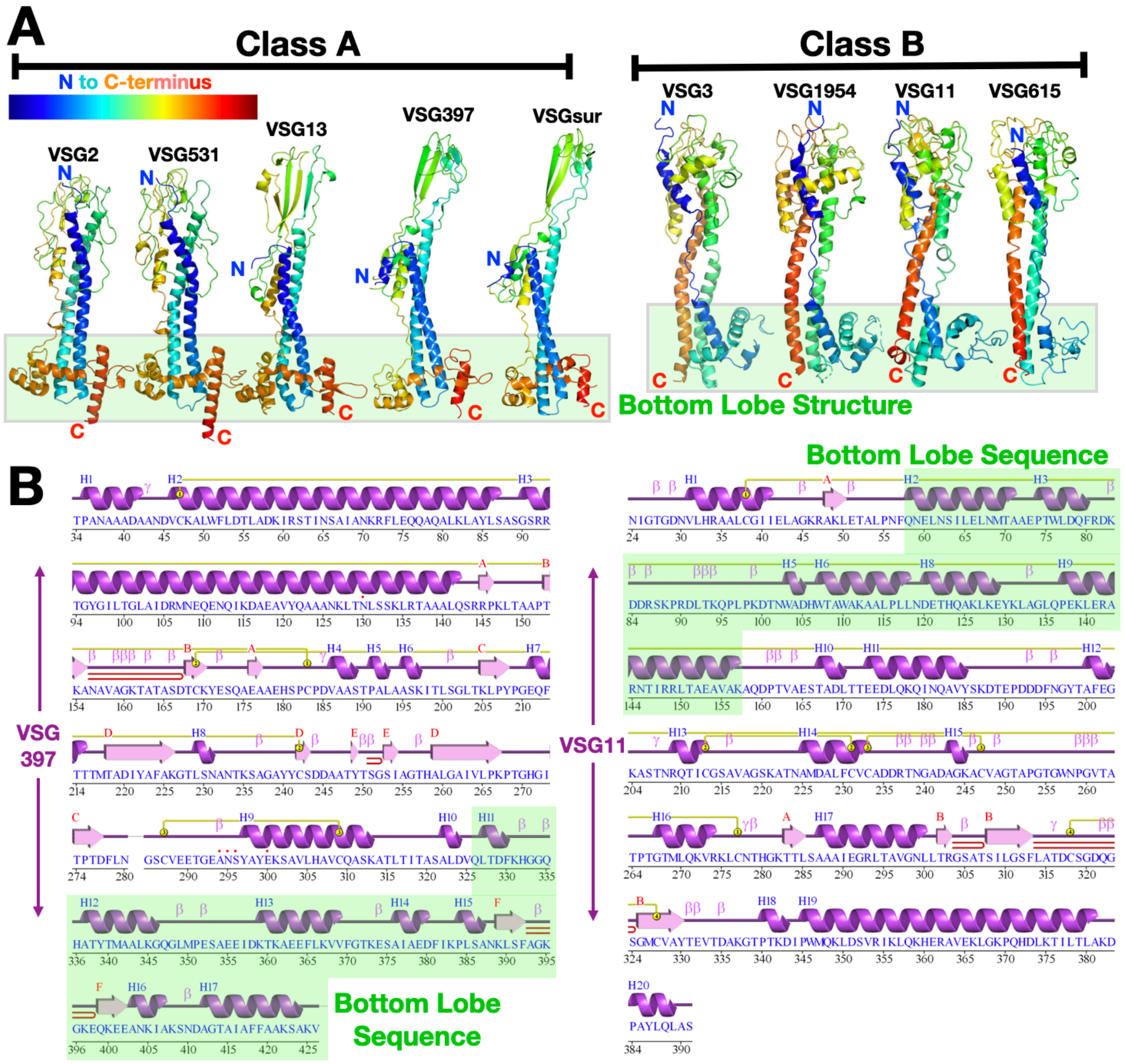


**Fig S4. Topology of Class A and Class B** (A) Two broad superfamily classes of VSGs identified through sequence analysis and defined by structural topology are shown here with representative structures. VSG monomers are shown as ribbon diagrams colored in a gradient from blue to red from N- to C-terminus. Structures reported in this manuscript are denoted in red and described in detail below. The lower, bottom lobe subdomains of the VSGs are highlighted in a green-tinted box. Structures drawn with CCP4*mg* (McNicholas et al. 2011). (B) Sequence and structurally-determined secondary structure of two representative VSGs from class A (VSG397) and class B (VSG11). The sequence region that forms the bottom lobe is indicated by green highlighting. Secondary structure illustrated with PDBSUM (Laskowski et al. 2018).

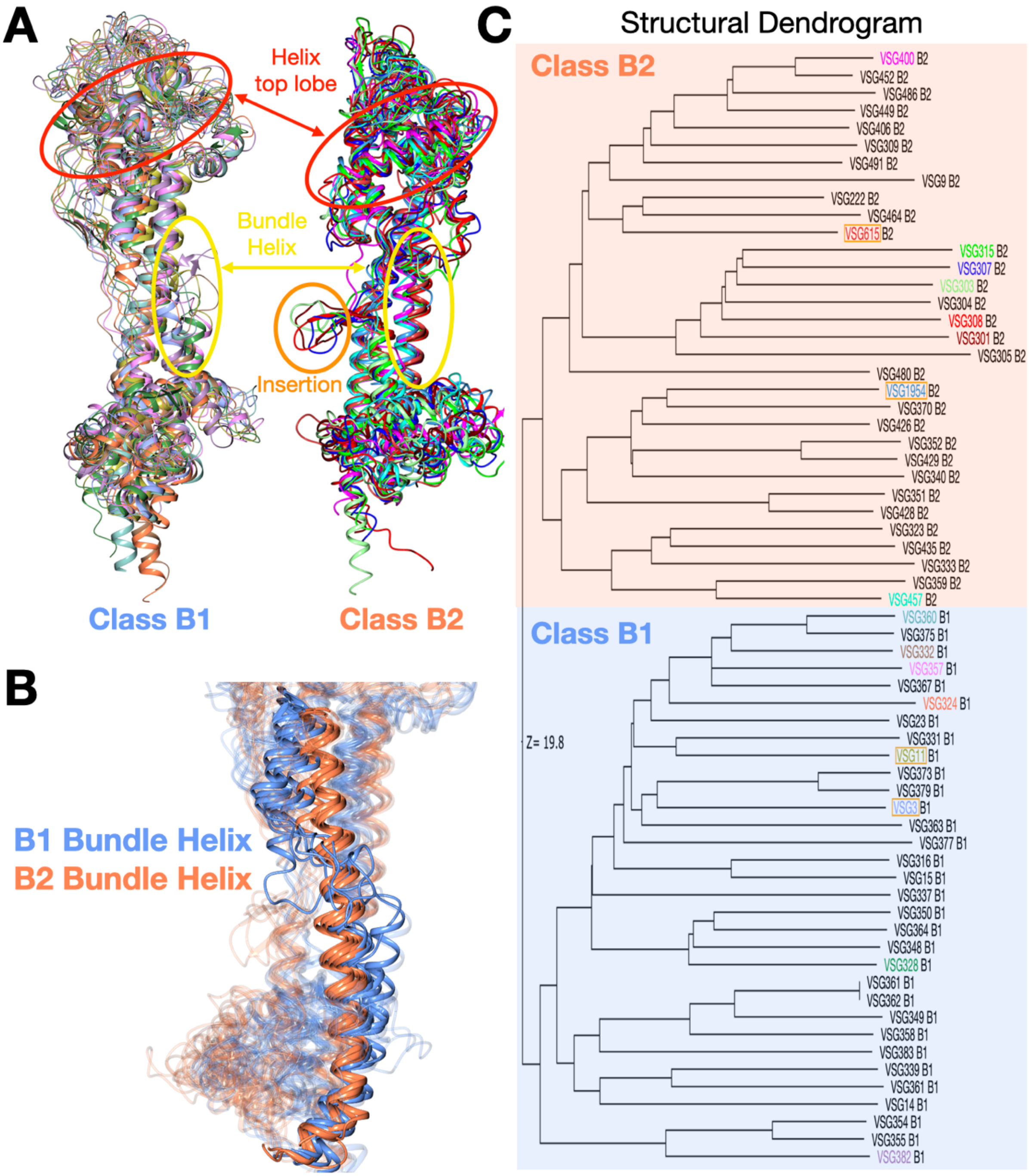


**Fig. S5. Subtle Structural Differences Between Class B1 and Class B2 VSGs. (A)** Alignments of class B1 and B2 VSGs (experimental and AlphaFold models), highlighting the top lobe helices in B2 but not in B1 (red circle) , the disordered bundle helix in B1 (yellow circle), and the inserted sequence in the bundle region (orange circle). The structural drawings match in color the names of the VSGs in panel (C). **(B)** Focus on the helixes in the 3-helix bundle differing between class B1 and B2. Orange is B2, showing that the helix is unbroken in most B2 VSGs. In B1 VSGs (light blue) the helix becomes disordered in the middle of the bundle and resumes a displaced helical structure afterward. **(C)** Structural dendrogram generated by the Dali Server comparing a number of experimental (VSG names with orange outline box) and AlphaFold models of the class B VSGs, showing that they split into two groups. B1 and B2 subgroups are shown with light blue and light orange backgrounds, respectively.

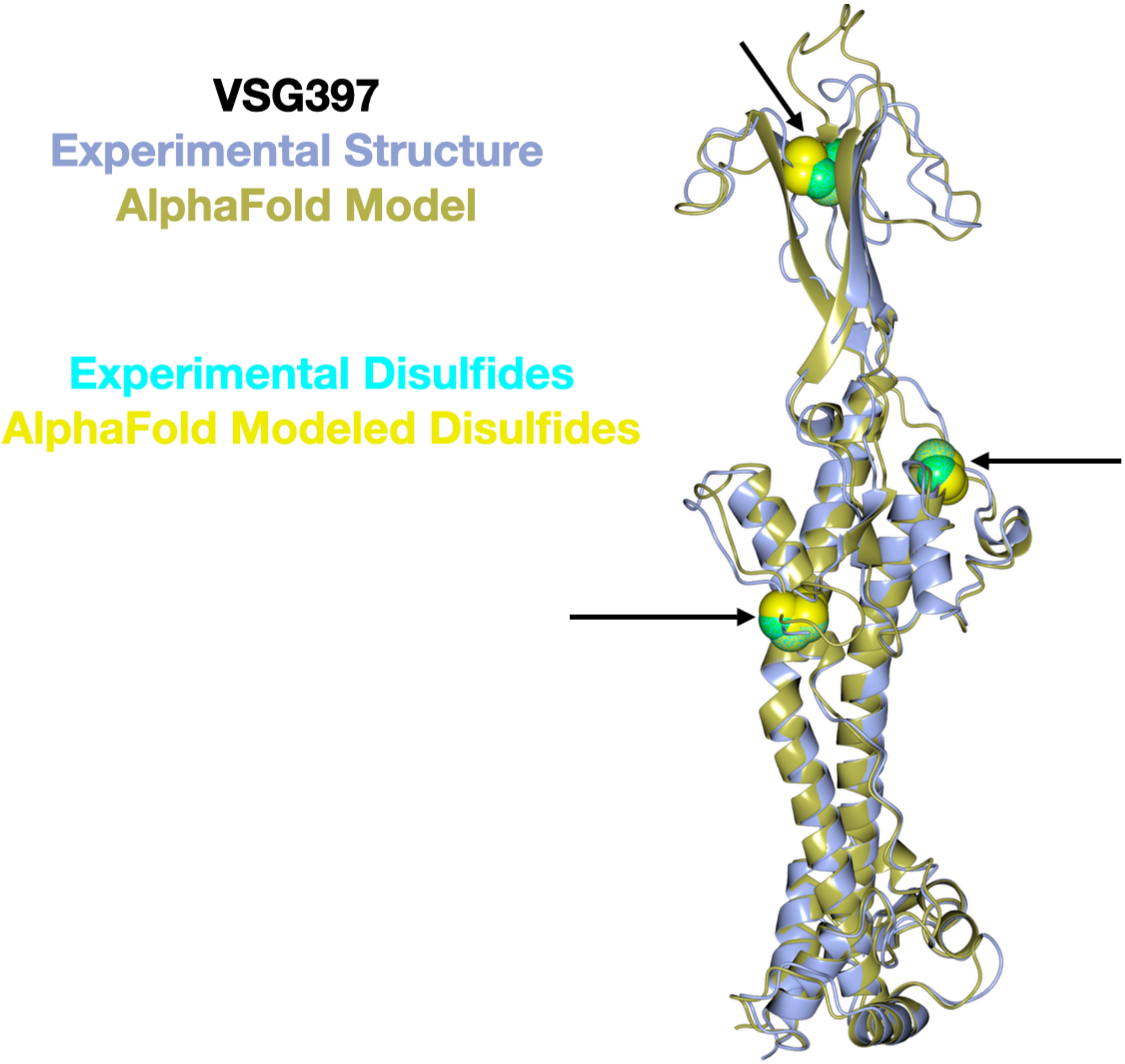


**Fig S6. Comparison of Experimental and AlphaFold Predicted Structures of VSG397.** Ribbon diagrams of the VSG397 monomers are shown in light blue and gold for the experimental and predicted structures, respectively. Disulfides are shown in cyan and yellow for the experimental and predicted structures, respectively, marked by black arrows.

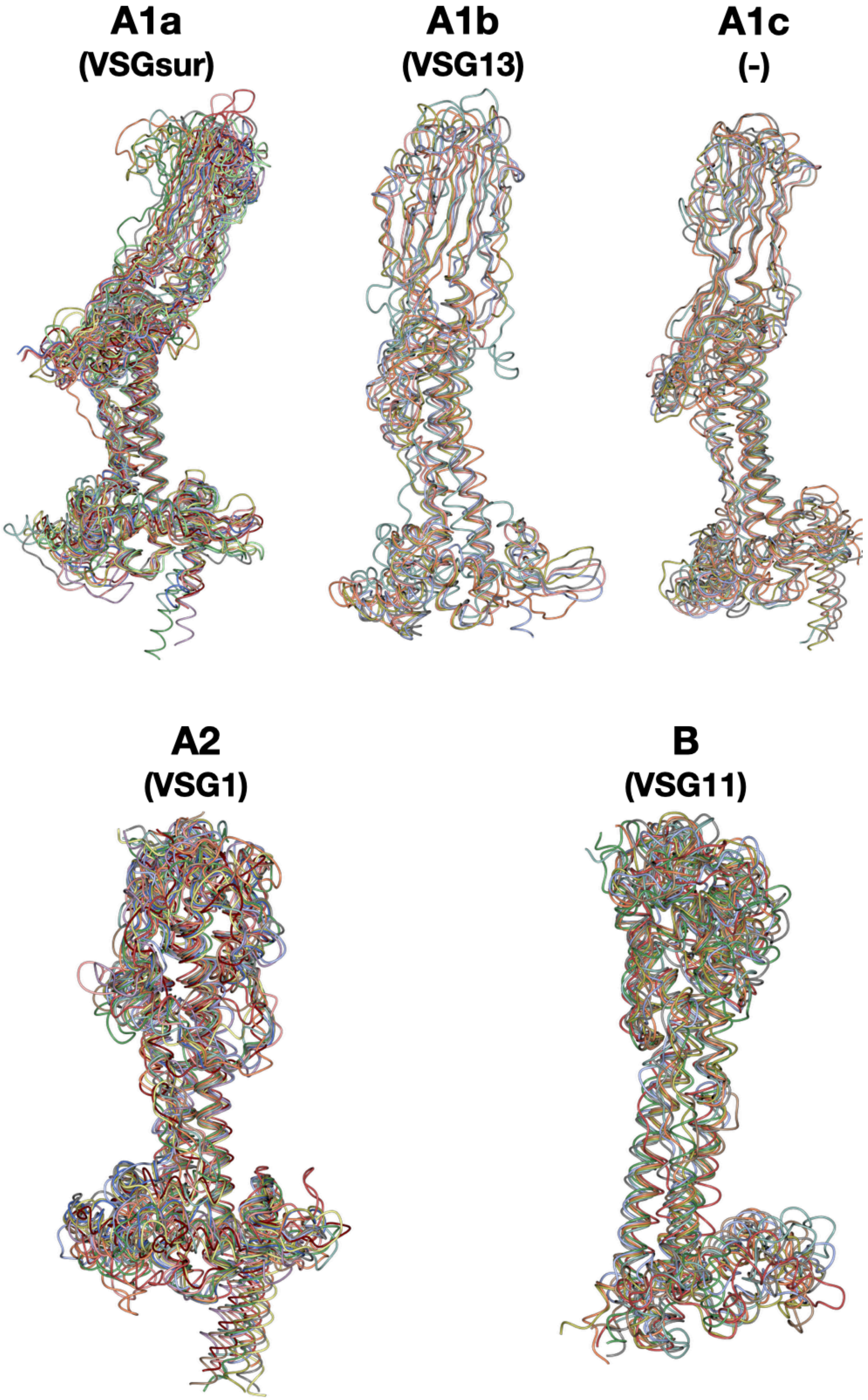


**Fig S7. AlphaFold Models of Sequences from VSG Structural Classes from *T. brucei brucei* 927.** Structures of aligned VSG monomers are drawn in different colors with a thin worm style. **(A)** Class A1a with models for the UniProtKB/Swiss-Prot accession numbers Q580P4, Q580P5, Q57TR8, Q57TR3, Q38CQ8, Q57X40, Q583L8, Q583L5, Q57X41, Q22KU2, Q38G13, Q380W0, and Q38CP5 aligned to VSGsur. **(B)** Class A1b with models for the UniProtKB/Swiss-Prot accession numbers Q57Y76, Q57X39, Q4GY52, Q586M4, Q57XI7 aligned to VSG13. **(C)** Class A1c with models for the UniProtKB/Swiss-Prot accession numbers Q380U5, Q380V8, Q380U9, Q380Y0, Q580N8, Q38CP2 aligned to an AlphaFold model for VSG21. **(D)** Class A2 with models for the UniProtKB/Swiss-Prot accession numbers Q57X38, Q580N9, Q57Z50, Q4FKU3, Q380X8, Q57XH3, Q38G16, Q57TR9, Q380W1, Q38CQ1, Q583L4, Q38G20 aligned to VSG1. **(E)** Class B with models for the UniProtKB/Swiss-Prot accession numbers Q380U8, Q583L3, Q4FKE9, Q38CP0, Q57TR6, Q387P0, Q57TR7, Q4GY50 aligned to VSG11. Alignments generated and illustrated with CCP4mg, each structure aligned in a different color.

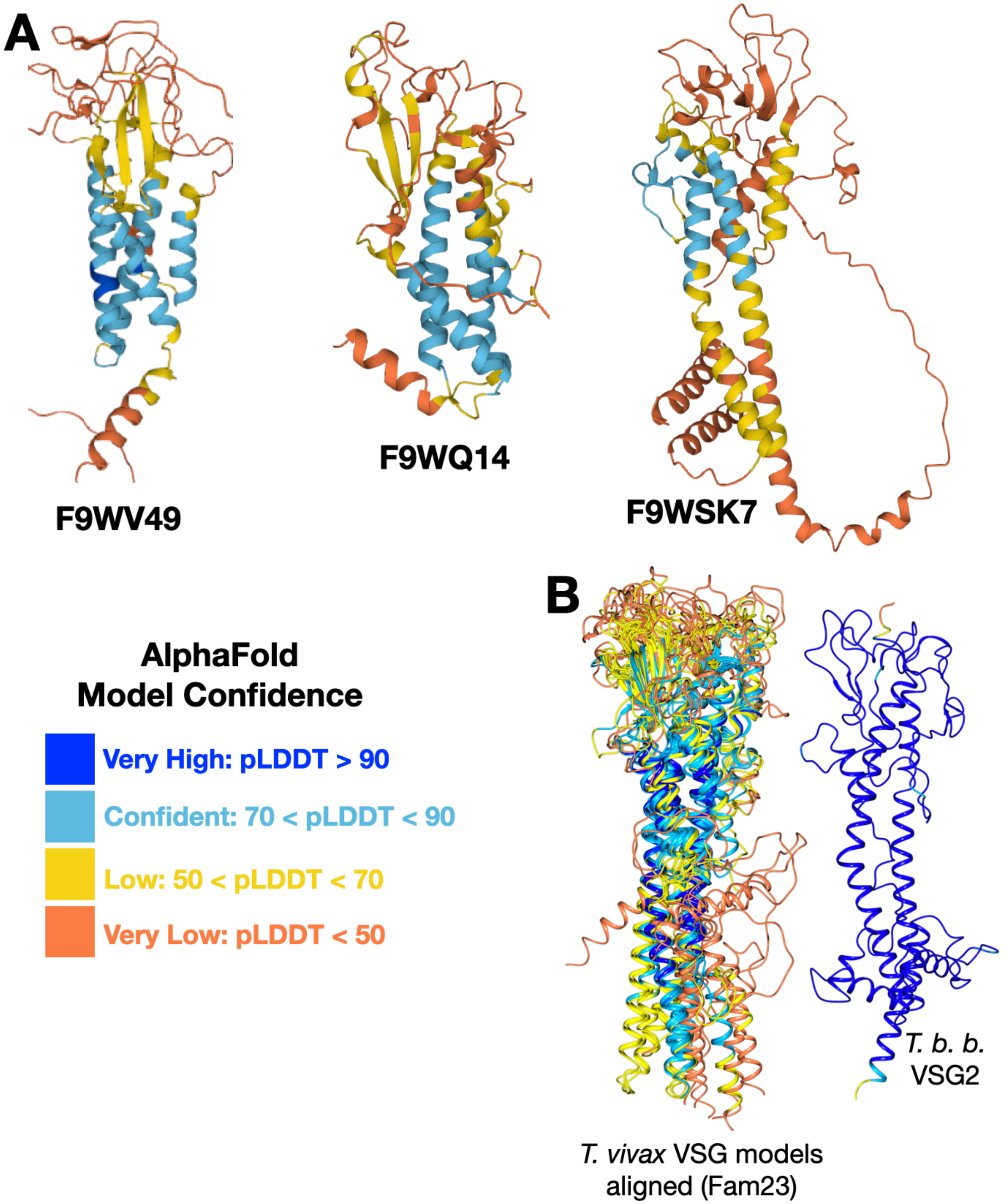


**Fig S8. AlphaFold *T. vivax* VSG Predictions.** (A) A collection of several *T. vivax* VSG protein structures predicted by AlphaFold and colored by pLDDT confidence score (as indicated). (B) Comparison of several *T. vivax* putative VSG structures (from Fam23 of the Fam23-26 VSG gene groups) and the prediction of *T. brucei brucei* VSG2, both colored by pLDDT confidence scores.

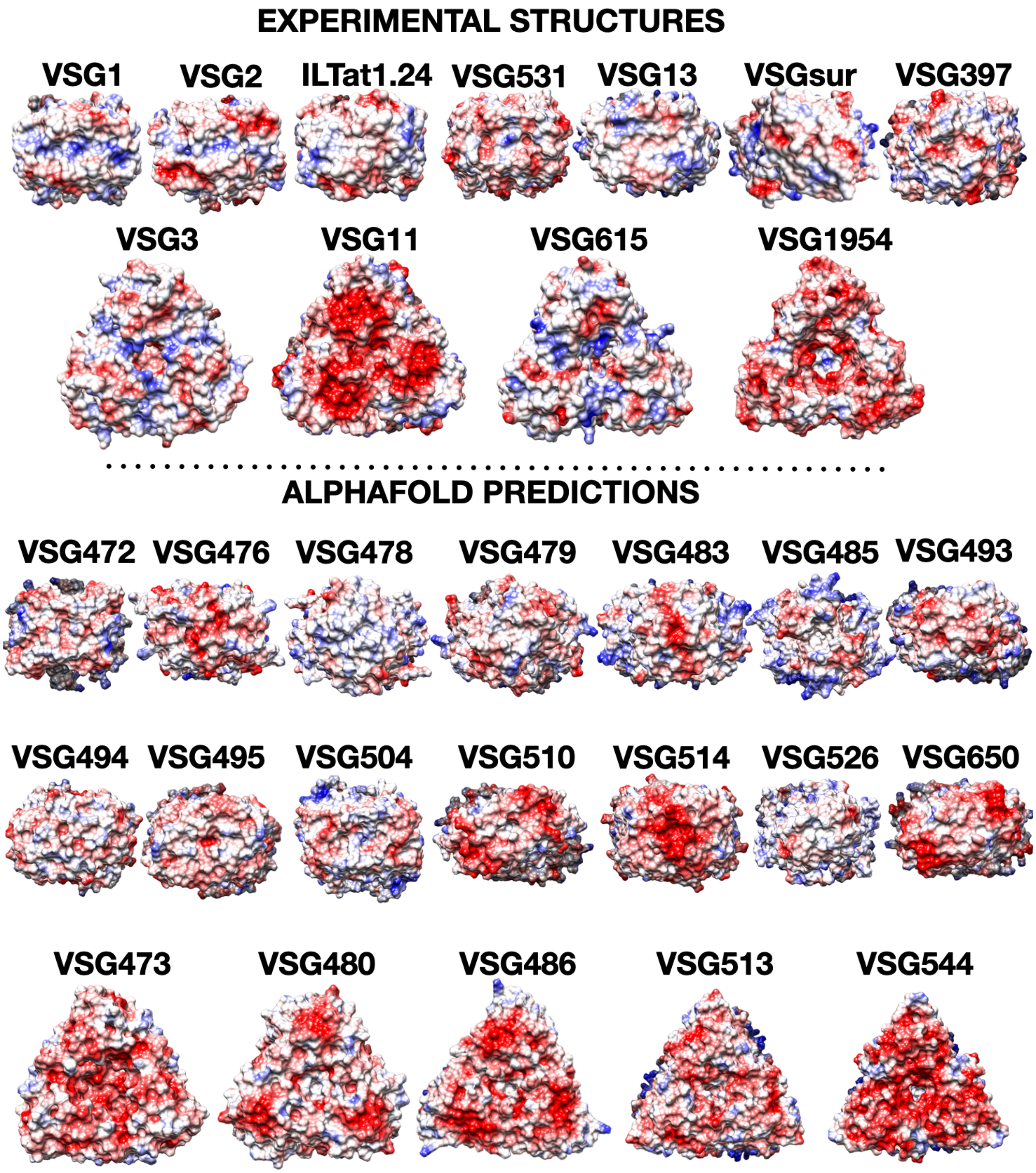


**Fig S9. VSG Molecular Surfaces.** Colored by charge distribution with blue indicating positive or basic, white neutral, and red acidic or negative (produced with Chimera(Pettersen et al. 2004) using the "Coulombic Surface Coloring" option). The top portion shows surfaces from experimental structures, the bottom the surfaces from a random assortment of predictions by AlphaFold (in either the dimeric or trimeric assemblies).

**Table S1: Crystallographic Statistics**

| Parameter | VSG11$_{WT}$-Iodine | VSG11$_{WT}$-Oil | VSG11$_{WT}$-AS | VSG11$_{N2C}$-18mer | VSG615 | VSG558 |
|---|---|---|---|---|---|---|
| Wavelength | 1.00 Å | 1.00 Å | 1.00 Å | 1.00 Å | 1.00 Å | 1.00 Å |
| Resolution range | 41.26-1.27 (1.32-1.27) | 35.28-1.23 (1.27-1.23) | 42.45-1.75 (1.81-1.75) | 48.81-2.59 (2.68-2.59) | 68.16-3.22 (3.34-3.22) | 46.17-1.96 (2.03-1.96) |
| Space group | P 3 2 1 | P 3 2 1 | P 3 2 1 | P 1 21 1 | P 1 21 1 | P 1 21 1 |
| Unit cell | 75.43 75.43 106.4 90 90 120 | 74.86 74.86 105.61 90 90 120 | 76.24 76.24 212.26 90 90 120 | 131.9 210.77 133.75 90 104.54 90 | 86.28 111.23 108.59 90 91.65 90 | 92.33 60.33 141.19 90 108.79 90 |
| Total reflections | 901512 (86366) | 938886 (70182) | 1384124 (124473) | 1484781 (140952) | 186820 (20643) | 548887 (49698) |
| Unique reflections | 92763 (9185) | 99721 (9827) | 72926 (7093) | 217669 (20848) | 30929 (3271) | 101158 (9338) |
| Multiplicity | 9.7 (9.4) | 9.4 (7.1) | 19.0 (17.5) | 6.8 (6.8) | 6.0 (6.3) | 5.4 (5.3) |
| Completeness (%) | 99.96 (100) | 99.91 (99.75) | 99.75 (97.77) | 99.42 (95.33) | 92.38 (97.84) | 95.08 (86.86) |
| Mean I/sigma(I) | 14.54 (1.27) | 15.79 (0.98) | 19.09 (0.98) | 14.56 (1.22) | 3.71 (0.63) | 5.66 (0.67) |
| Wilson B-factor | 14.58 | 13.37 | 32.69 | 66.92 | 85.12 | 35.08 |
| R-merge | 0.0903 (1.683) | 0.0754 (1.777) | 0.09623 (2.514) | 0.09065 (1.294) | 0.3959 (2.754) | 0.1587 (2.803) |
| R-meas | 0.0954 (1.781) | 0.0796 (1.918) | 0.09892 (2.588) | 0.0983 (1.402) | 0.4333 (2.998) | 0.1757 (3.104) |
| R-pim | 0.0304 (0.5777) | 0.0258 (0.7099) | 0.02264( 0.6037) | 0.03766( 0.5325) | 0.1736( 1.171) | 0.07463 (1.319) |
| CC1/2 | 0.999 (0.471) | 0.999 (0.388) | 1 (0.348) | 0.999 (0.545) | 0.978 (0.127) | 0.996 (0.254) |
| CC* | 1 (0.8) | 1 (0.748) | 1 (0.718) | 1 (0.84) | 0.995 (0.474) | 0.999 (0.636) |
| Reflections (refinement) | 92761 (9185) | 99650 (9807) | 72898 (7073) | 217644 (20848) | 30849 (3257) | 100878 (9148) |
| Reflections (R-free) | 4639 (460) | 4979 (489) | 3645 (355) | 10882 (1043) | 1525 (171) | 5031 (441) |
| R-work | 0.2063 (0.2937) | 0.1614 (0.2734) | 0.1991 (0.3171) | 0.2168 (0.3416) | 0.2828 (0.3764) | 0.2129 (0.3407) |
| R-free | 0.2300 (0.29927) | 0.1863 (0.3119) | 0.2378 (0.3486) | 0.2673 (0.3695) | 0.3210 (0.4080) | 0.2505 (0.3565) |
| CC(work) | 0.943 (0.692) | 0.964 (0.657) | 0.957 (0.609) | 0.933 (0.670) | 0.919 (0.290) | 0.958 (0.601) |
| CC(free) | 0.934 (0.680) | 0.960 (0.503) | 0.957 (0.777) | 0.910 (0.581) | 0.865 (0.137) | 0.956 (0.532) |
| all atoms* | 3338 | 3317 | 6295 | 43923 | 13915 | 12172 |
| Protein atoms | 2782 | 2822 | 5604 | 42731 | 13713 | 10987 |
| ligands | 175 | 154 | 240 | 909 | 194 | 546 |
| solvent | 452 | 412 | 451 | 283 | 8 | 639 |
| Protein residues | 366 | 368 | 736 | 5835 | 1872 | 1443 |
| RMS(bonds) | 0.012 | 0.004 | 0.008 | 0.005 | 0.003 | 0.008 |
| RMS(angles) | 1.24 | 0.83 | 1.06 | 0.76 | 0.6 | 1.06 |
| Ramachandran favored (%) | 97.25 | 97.81 | 96.31 | 91.86 | 83.06 | 95.96 |
| Ramachandran allowed (%) | 2.75 | 2.19 | 3.55 | 7.20 | 14.74 | 3.76 |
| Ramachandran outliers (%) | 0.00 | 0.00 | 0.14 | 0.94 | 2.21 | 0.28 |
| Rotamer outliers (%) | 0.70 | 0.33 | 2.86 | 3.93 | 1.41 | 1.40 |
| Clashscore | 2.11 | 1.73 | 2.94 | 8.55 | 12.02 | 6.07 |
| Average B | 20.70 | 20.73 | 42.79 | 95.05 | 85.30 | 45.46 |
| protein | 19.77 | 19.21 | 42.38 | 94.89 | 85.06 | 45.11 |
| ligands | 21.35 | 24.55 | 50.80 | 113.48 | 104.80 | 54.27 |
| solvent | 26.29 | 30.43 | 43.61 | 59.82 | 20.56 | 43.91 |
| TLS groups | 6 | | 15 | 118 | | 24 |

Statistics for the highest-resolution shell are shown in parentheses.

*All atoms refers to non-hydrogen atoms
